# Large-scale Proteomic and Phosphoproteomic Analysis of Maize Seedling Leaves During De-etiolation

**DOI:** 10.1101/2020.03.13.977843

**Authors:** Zhi-Fang Gao, Zhuo Shen, Qing Chao, Zhen Yan, Xuan-Liang Ge, Tiancong Lu, Haiyan Zheng, Chun-Rong Qian, Bai-Chen Wang

## Abstract

De-etiolation consists of a series of developmental and physiological changes that a plant undergoes in response to light. During this process light, an important environmental signal, triggers the inhibition of mesocotyl elongation and the production of photosynthetically active chloroplasts, and etiolated leaves transition from the “sink” stage to the “source” stage. De-etiolation has been extensively studied in maize (*Zea mays L*). However, little is known about how this transition is regulated. In this study, we describe a quantitative proteomic and phosphoproteomic atlas of the de-etiolation process in maize. We identified 16,420 proteins and quantified 14,168. In addition, 8,746 phosphorylation sites within 3,110 proteins were identified. From the proteomic and phosphoproteomic data combined, we identified a total of 17,436 proteins, 27.6% of which are annotated protein coding genes in the *Zea_mays* AGPv3.28 database. Only 6% of proteins significantly changed in abundance during de-etiolation. In contrast, the phosphorylation levels of more than 25% of phosphoproteins significantly changed; these included proteins involved in gene expression and homeostatic pathways and rate-limiting enzymes involved in photosynthesis light and carbon reactions. Based on phosphoproteomic analysis, 34% (1,057) of all phosphoproteins identified in this study contained more than three phosphorylation sites, and 37 proteins contained more than 16 phosphorylation sites, which shows that multi-phosphorylation is ubiquitous during the de-etiolation process. Our results suggest that plants might preferentially regulate the level of PTMs rather than protein abundance for adapting to changing environments. The study of PTMs could thus better reveal the regulation of de-etiolation.

## Introduction

Proteotype is the proteomic state of a cell, and it reflects the integration of a cell’s genotype, developmental history and environment [1]. Diversity of proteotype in a cell or tissue mainly comes from two forms: variants affecting the primary amino acid sequence and posttranslational modifications (PTMs) [2]. Although the genotype specifies the potential phenotype of an organism, proteins that implement cellular processes, and the interactions between these proteins and outside environment, dictate the actual phenotype. Therefore, to fully understand the biology of an organism and its constituent parts, the knowledge of proteotype, or protein complement is required.

PTMs of the protein is an important component of the proteotype, which may affect protein functions, such as protein phosphorylation. Generally speaking, a newly synthesized protein may not have a biological function until it is modified [3]. PTMs provide a more precise and elegant mechanism to control cellular function than regulation of gene expression [4]. For instance, PINFORMED1 (PIN1) shows a tissue-specific difference in phosphorylation in the maize leaf that correlates with changes in polarized localization of PIN1 in epidermal cells during development [5]. In PTM databases, more than 300 different types of PTMs have been described and the number is still increasing [6].

Protein phosphorylation is an important and major type of PTM, which has been extensively studied since it was first reported in 1926 [7]. According to published data, protein phosphorylation is one of the most abundant modifications in all biological species, representing 53.5% of all PTMs [8]. The conversion between phosphorylation and dephosphorylation of specific sites can alter the molecular conformation of the protein, potentially affecting enzyme activity, substrate specificity, structural stability or intracellular localization, and thus the regulation of biological processes [8, 9].

Many proteins contain multiple phosphosites. On one hand, different phosphosites can regulate different functions of the target protein. For example, phosphorylation of proteins at different sites activates or inhibits their activities [10]. On the other hand, a combination of multiple phosphosites that have similar functions in the same protein may amplify the effect of phosphorylation. Moreover, phosphorylation of multiple sites on the same protein can function as a molecular switch that allows biological crosstalk between different redundant and alternative pathways [11]. Phosphoproteome analysis, which includes phosphoprotein identification, the exact mapping of phosphorylation sites, quantification of phosphorylation, and identification the associated biological functions, is an effective approach for analyzing these biological regulatory networks at a global level [9, 12].

Seedling de-etiolation is a complex but precisely regulated process. During subterranean growth, dark-grown or etiolated seedlings have fast-growing hypocotyls (dicotyledonous plants) or mesocotyls (monocotyledonous plants) that allow them to rapidly reach the light, together with a protective apical hook and appressed cotyledons (dicotyledonous plants) or a protective coleoptile (monocotyledonous plants) with undeveloped chloroplasts. At the soil surface, incident light represses hypocotyl or mesocotyl elongation and stimulates cotyledon separation (dicots) or leaf expansion (monocots), congruent with the development of functional chloroplasts, thus enabling light capture for photosynthesis [13, 14]. Several key regulators of the de-etiolation process have been identified, including Constitutive Photomorphogenic 1 (COP1), Elongated Hypocotyl 5 (HY5), and Phytochrome-interacting Factors (PIFs), which play essential roles in regulating the massive reprogramming of the plant transcriptome during de-etiolation [15−17]. Moreover, phosphorylation modification plays an essential role in the regulation of these key regulators. For example, PHYs are unphosphorylated and located in the cytosol in the dark. After illumination with light, they are converted to active Pfr forms and phosphorylated, and then rapidly localize to the nucleus where they phosphorylate downstream proteins, such as PIFs. Phosphorylated PIFs are targeted to the proteasome and degraded, resulting in the promotion of photomorphogenesis [17, 18].

Previous studies indicated that a significant portion of the genome, at least 20% in both *Arabidopsis* and rice, is differentially expressed between seedlings that are undergoing photomorphogenesis and those that are undergoing skotomorphogenesis [19, 20]. However, it has become clear that mRNA levels are poorly correlated with protein abundance [1, 21]. To bridge this gap, proteomics studies have been performed on *Arabidopsis* [22], rice [23, 24], and maize [25] seedlings undergoing de-etiolation. However, due to the limited ability to identify and quantify protein phosphorylation using the proteomic methods available at the time these studies were performed, only several dozen proteins were found to have differences in protein phosphorylation level. Therefore, a much deeper proteomic survey is needed to reveal the mechanism by which phosphorylation regulates seedling de-etiolation.

Besides being the world’s largest crop in terms of production, maize is also an important model plant for basic research, especially as a C4 model plant for photosynthesis research. The completion of the B73 maize genome [26] has facilitated the use of large-scale transcriptome and proteome data to reveal the mechanisms underlying various maize developmental and physiological processes. For example, researchers have created large data resources for C4 photosynthesis research, including complementary RNA-seq [27], proteomics [28, 29], and phosphoproteomics [30, 31] data for a developmental gradient of the maize leaf. In these studies, the mRNA and protein contents at successive stages of photosynthetic development were analyzed. However, our understanding of how the proteome changes during a given developmental process of maize is still incomplete. In the present study, we performed 3D-HPLC-MS/MS and 2D-HPLC-MS/MS to obtain deep proteomic information for maize seedlings undergoing de-etiolation, and used two methods, IMAC (Immobilized Metal ion Affinity Chromatography) and TiO_2_, to enrich phosphopeptides to obtain deep phosphoproteomic information. Our results provide abundant data for better understanding the regulation of de-etiolation in maize.

## Results and Discussion

### Strategy for quantitative analysis of the maize leaf proteome and phosphoproteome

To perform a deep analysis of the maize proteome and phosphoproteome during de-etiolation, the first leaves of etiolated 7-day-old maize seedlings (B73 inbred) were illuminated with white light and harvested at 0 h, 1 h, 6 h and 12 h for mass spectrometry analysis (**Figure 1**). We chose these samples for analysis because in our previous studies [25, 31, 32] we found that the first leaves of 7-day-old etiolated maize seedlings showed no significant changes in growth compared with light-grown seedlings when subjected to illumination for 12 hours, and few proteins involved in development and growth were found to be differently expressed when comparing the proteomes of etiolated and normal leaves. Moreover, the etiolated leaves turned green after 12 hours of illumination, which indicated that processes related to greening in leaves were completed. Total protein was extracted, and each sample was labeled with one of four iTRAQ tags and combined after trypsin digestion (Figure 1A). To obtain deep proteomic information, the iTRAQ-labeled peptides were analyzed with 3D-HPLC-MS/MS. Specifically, the labeled peptides were first separated into six primary fractions by strong cation exchange chromatography (SCX), and each primary fraction was then separated into 14 secondary fractions using high pH reversed-phase chromatography (HpH RP) (Figure 1B). Finally, the resulting 84 fractions were analyzed with HPLC-MS/MS. The spectra were searched against maize annotated proteins downloaded from EnsEMBL Plants release 29 [33] using the X!Tandem search algorithm and filtered at 1% FDR (Figure 1D). In total, 15,206 proteins were identified. Of these proteins, 13,115 were quantified with at least one uniquely mapped peptide and at least two quantifiable spectra (**Figure 2A**, Supplementary Table 1). To validate this result, the maize leaf proteome was reanalyzed in two independent experiments using 2D-HPLC-MS/MS, in which iTRAQ-labeled peptides were separated into 17 fractions using HpH RP and then analyzed with HPLC-MS/MS (Figure 1B). Using the same criteria mentioned above, 11,129 proteins were identified and 9,933 proteins were quantified using 2D-HPLC-MS/MS (Figure 2A, Supplementary Table 1). The Pearson correlation coefficients determined by comparing the protein abundances from 3D proteome analysis with those from replicates 1 and 2 of the 2D proteome analysis were 0.76 and 0.79, respectively, demonstrating that there was good correlation between the two methods (Supplementary Figure 1). A total of 9,915 proteins were identified with both 2D- and 3D-HPLC-MS/MS, and of these proteins, 8,880 were quantified (Figure 2A). Moreover, the dynamic changes of 10 quantified proteins were confirmed by western blot analysis (Figure 2D); the accumulation patterns of these proteins were consistent with the patterns determined from HPLC-MS/MS data (Figure 2C, Supplementary Table 1). Taken together, using the two approaches, we identified a total of 16,420 proteins encoded by 15,653 genes, and of these proteins, 14,168 encoded by 13,613 genes were quantified.

**Figure 1.**
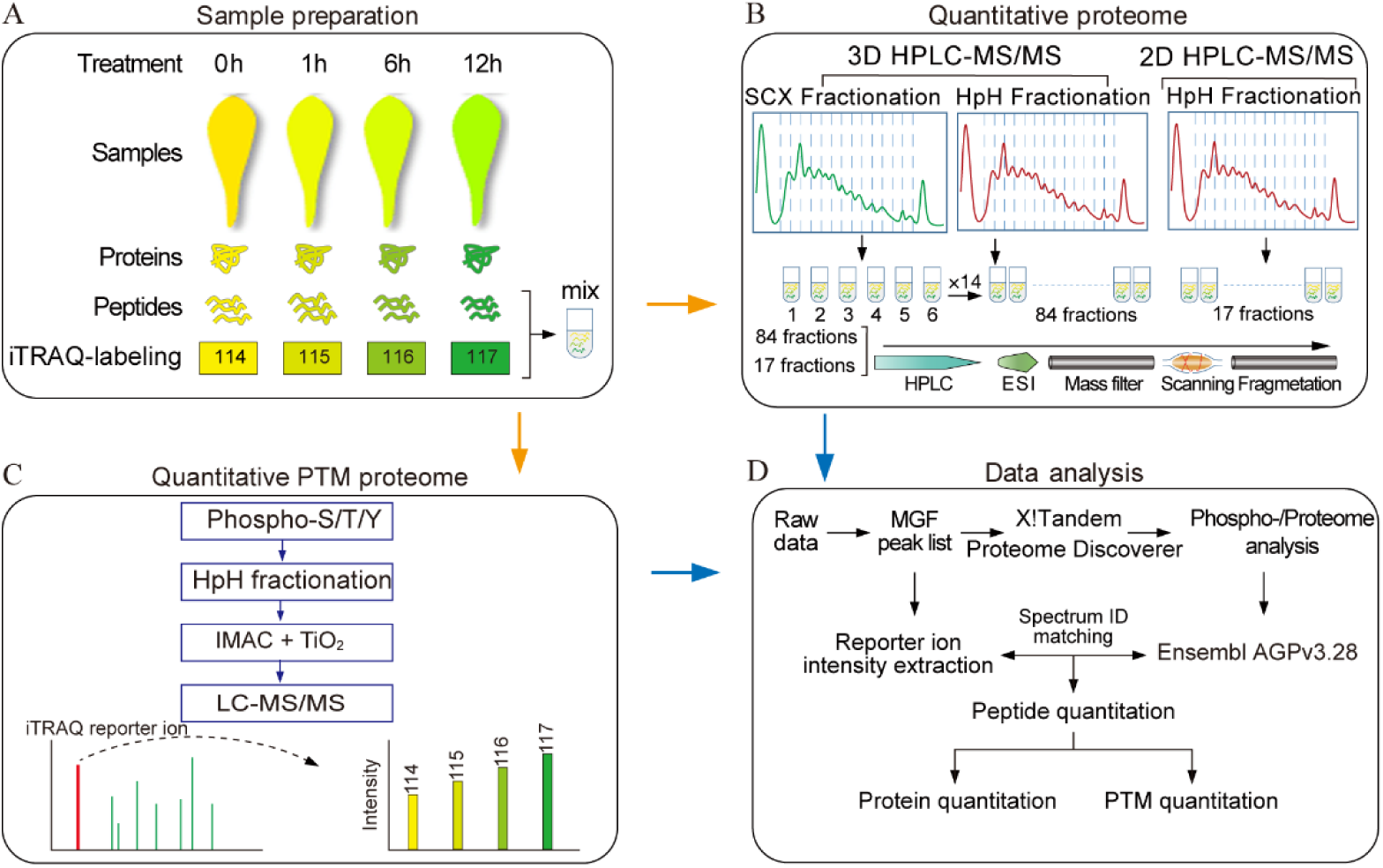
Experimental workflow for the proteome and phosphoproteome analyses of de-etiolated maize seedling leaves. **A**. Etiolated maize seedlings (B73 inbred) were illuminated with white light, and the first leaves were harvested after 0 h, 1 h, 6 h and 12 h. Total protein was extracted, and after trypsin digestion, proteins from each sample were labeled with one of four iTRAQ tags and combined. **B**. The analysis of iTRAQ-labeled peptides with 3D-HPLC-MS/MS and 2D-HPLC-MS/MS. In 3D-HPLC-MS/MS analysis, the labeled peptides were first separated into six fractions by strong cation exchange chromatography (SCX) then separated into 14 secondary fractions using high pH reversed-phase chromatography (HpH-RP). In 2D-HPLC-MS/MS analysis, iTRAQ-labeled peptides were separated into 17 fractions by HpH chromatography before being subjected to HPLC-MS/MS. ESI: Electron Spray Ionization **C**. Enrichment and purification of phosphorylated peptides. Phospho-S/T/Y: peptides containing phosphosites on Ser/Thr/Tyr; IMAC: Immobilized Metal ion Affinity Chromatography; TiO_2_: Titanium dioxide. **D**. Spectra were searched against annotated databases. MGF: Mascot Generic Format.

**Figure 2.**
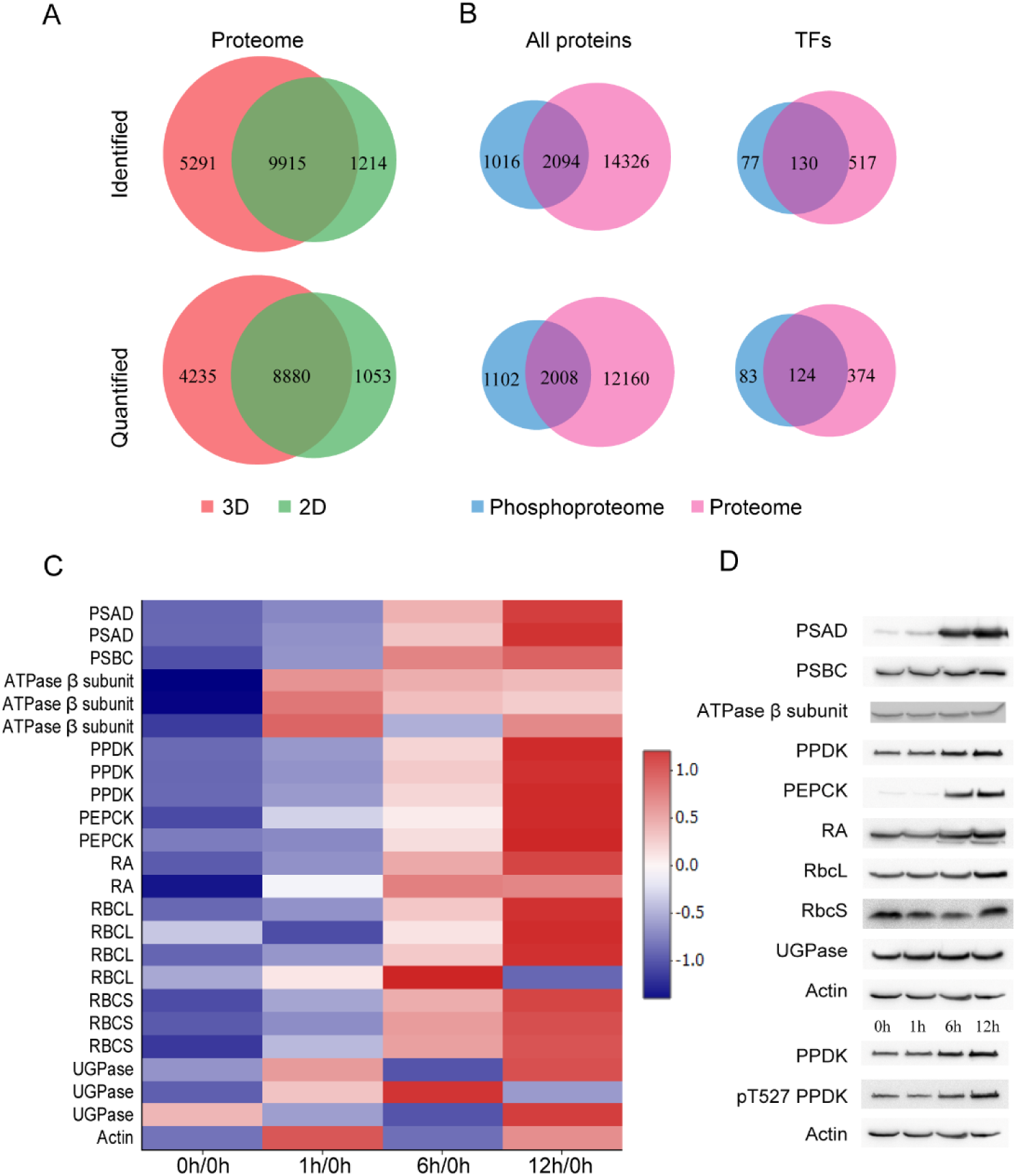
Overview of the number of proteins and phosphoproteins identified in de-etiolated maize seedling leaves. **A**. Venn diagram showing the overlap in the number of proteins identified (top) and quantified (bottom) in de-etiolated maize seedling leaves using 2D- and 3D-HPLC-MS/MS. **B**. Venn diagrams showing the overlap in the number of proteins (left) and the number of TFs (right) identified (top) and quantified (bottom) in the proteomic and phosphoproteomic analyses of de-etiolated maize seedling leaves. In the phosphoproteomic analysis, 3,110 phosphoproteins were quantified, 2,008 of which matched quantified proteins. A total of 647 and 203 TFs were identified among the proteins and phosphoproteins, respectively. **C**. Heat map illustrating the dynamic changes in expression of 10 quantified proteins. The blue color represents low abundance while the red color represents high abundance. **D**. Western blot analysis of the 10 quantified proteins shown in C. PSAD: AD subunit of the PSII complex; PSBC: BC subunit of the PSII complex; ATPase β subunit: the β subunit of ATPase; PPDK: Pyruvate orthophosphate dikinase; PEPCK: Phosphoenolpyruvate carboxykinase; RA: Rubisco activase; RbcL: Large subunit of Rubisco; RbcS: small subunit of Rubisco; UGPase: UDP-glucose pyrophosphorylase; pT527 PPDK: PPDK phosphorylated on Thr527.

Characterization of myriad PTMs is another key aspect of proteome profiling. Here, we exhaustively studied one important PTM event, serine/threonine/tyrosine phosphorylation (pS/pT/pY), in de-etiolated maize seedling leaves. Phosphorylated peptides were enriched using IMAC (Immobilized Metal ion Affinity Chromatography) and TiO_2_ in parallel, and these peptides were then combined for MS analysis. The spectra were searched against the *Zea mays* database using MASCOT or MSAmanda in the Proteome Discoverer environment (Figure 1C). By using stringent cutoff criteria (see Materials and Methods), phosphorylation on 9,528 S/T/Y residues (sites) representing 3,110 proteins was quantified (Figure 2B, Supplementary Table 2). We identified 1,102 phosphorylated proteins that did not overlap with the quantified proteins, even those from 3D-HPLC-MS/MS, suggesting that most are low abundant proteins that could not be identified without enrichment (Figure 2B, Supplementary Table 3). To prevent possible biases due to variation in protein expression, the relative intensities of the phosphopeptides were normalized against changes in protein abundance [34]. Finally, 2,008 proteins with normalized phosphorylation levels (NPLs) were used for further analysis of phosphorylation dynamics.

Integrating the proteome and phosphoproteome results, we identified a total of 17,436 proteins encoded by 15,970 genes, including 721 (663 genes) transcription factors (TFs) (Figure 2, Supplementary Table 4).

### Dynamic reprogramming of the maize leaf proteome

To better understand the molecular mechanism of maize seedling photomorphogenesis, we systematically investigated the proteome dynamics during de-etiolation. We used a strict cutoff criterion, fold-change in abundance ≥1.5 or ≤0.67, to identify proteins with significant changes in abundance, i.e. differentially expressed proteins (DEPs) [35]. To our surprise, only 998 (7.04%) of the 14,168 quantified proteins, representing 980 genes, significantly changed in abundance during the de-etiolation process (Supplemental Table 1). To reveal patterns in the changes in DEP accumulation during de-etiolation, we firstly performed hierarchical clustering analyses using the average fold change in intensity ratios. As shown in **Figure 3A**, DEPs were divided into four clusters based on accumulation pattern, and nearly half of the proteins were assigned to cluster 2, in which the protein abundance continuously increased with prolonged illumination. Conversely, the abundances of proteins belonging to cluster 4 were dramatically downregulated after illumination. We next performed GO enrichment analysis of all DEPs (Figure 3B). We found that biological pathway (BP) terms for photosynthesis, homeostatic process, response to cold, generation of precursor metabolites and energy, DNA replication initiation, G-protein coupled receptor protein signaling pathway and apoptosis were highly enriched in DEPs. Of 52 proteins involved in photosynthesis, the abundance of only 6 (11.54%) significantly increased after 1 h, while 45 (86.54%) and 50 (96.15%) increased in abundance after 6 h and 12 h, suggesting that establishment of the photosynthetic machinery mainly occurred after 6 hours of illumination. We also performed GO enrichment analysis for DEPs belonging to cluster 2 and cluster 4 (**Figure 4**). Four pathways were highly enriched in cluster 2 proteins, which continuously increased in abundance after illumination: response to freezing, photosynthesis, homeostatic process and generation of precursor metabolites and energy.

**Figure 3.**
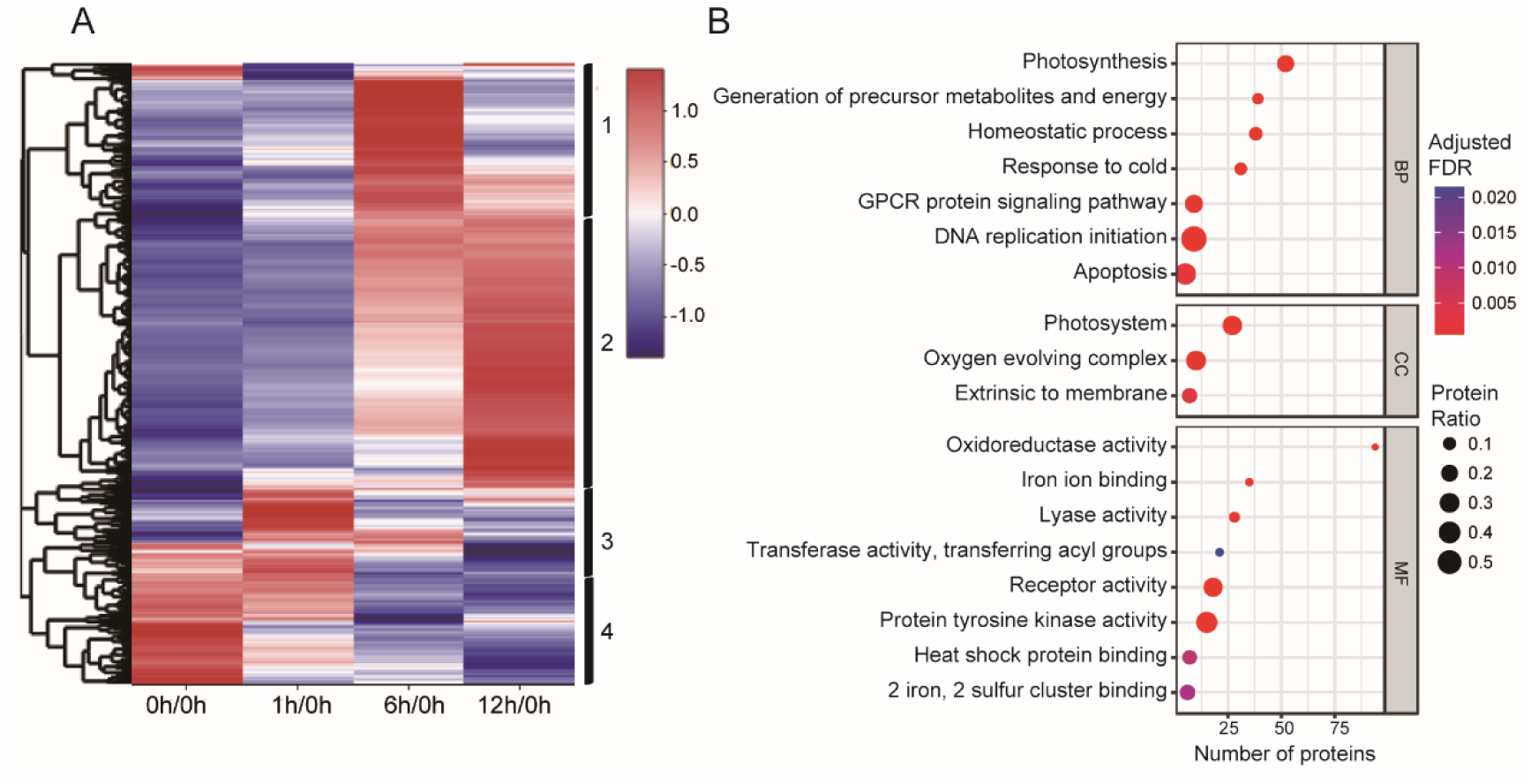
Proteome dynamics. **A**. Heat map of differentially expressed proteins (DEPs). The blue color represents low abundance while the red color represents high abundance. Vertical black lines on the right indicate the four clusters defined based on expression pattern. **B**. GO enrichment analysis of the DEPs identified in this study. Based on GO slim terms all DEPs were assigned to biological process (BP), cellular component (CC) and molecular function (MF) GO categories. Terms that were significantly enriched in DEPs (adjusted FDR≤ 0.05) are shown. The protein ratio is the ratio of the number of DEPs annotated to a certain GO term (adjusted FDR≤ 0.05) to the total number of proteins in the B73 maize genome assigned to that term. The horizontal axis indicates the total number of DEPs annotated to each GO term. GPCR protein signaling pathway: G-protein coupled receptor protein signaling pathway.

**Figure 4.**
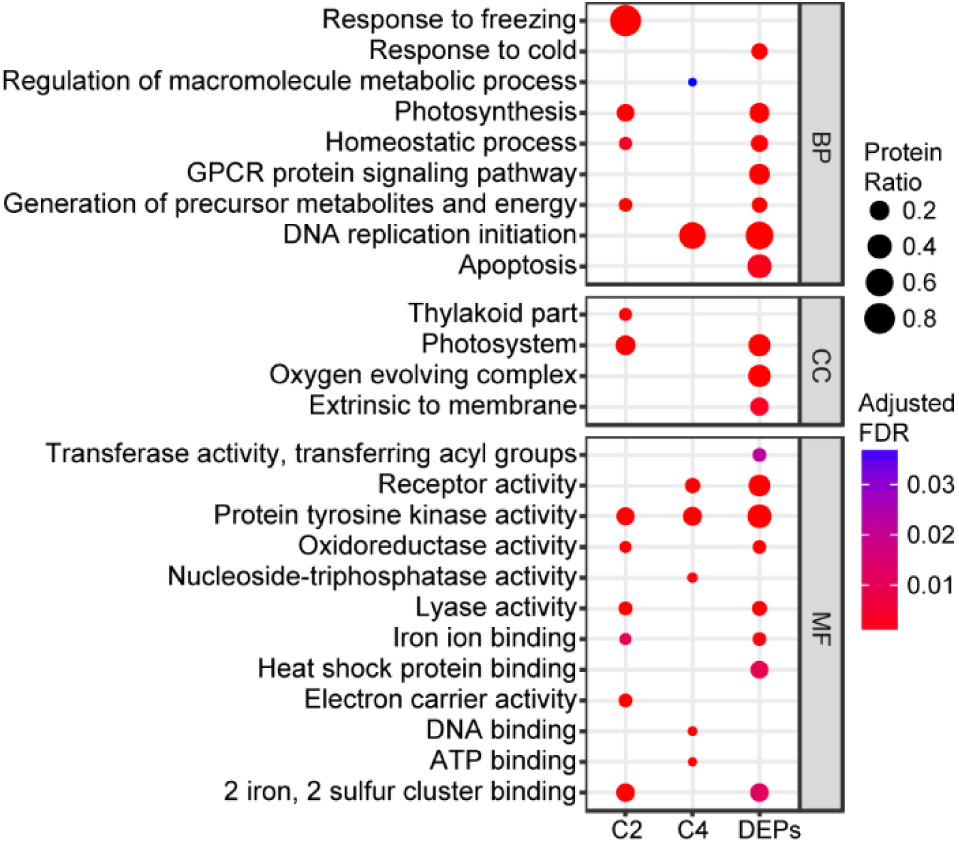
The enrichment of GO categories in DEPs belonging to cluster 2 and cluster 4 Based on GO slim terms, the DEPs belonging to cluster 2 (continuously upregulated during illumination) and cluster 4 (dramatically downregulated upon illumination) were assigned to biological process (BP), cellular component (CC) and molecular function (MF) GO categories. Terms that were significantly enriched in DEPs and proteins belonging to cluster 2 and cluster 4 (adjusted FDR≤ 0.05) are shown. The protein ratio is the ratio of the number of DEPs annotated to a certain GO term (adjusted FDR≤ 0.05) to the total number of proteins in the B73 maize genome assigned to that term. GPCR protein signaling pathway: G-protein coupled receptor protein signaling pathway; C2: Cluster 2; C4: Cluster 4; DEPs: differentially expressed proteins.

Numerous studies have shown that there is a complex cross-talk between pathways in response to light and low temperature although the mechanism remains poorly understood. For example, PIF3 and HY5 are key regulators in light response, besides they both play vital roles in response to low temperature in *Arabidopsis* [36−38]. In present study, when etiolated maize undergone photoporphogenesis, lots of proteins involving in response to light signals were changed in abundance, which night also play roles in resisting cold stress, so terms of response to freezing was enriched in cluster 2 proteins. In contrast, DNA replication initiation and regulation of macromolecule metabolic process were the most highly enriched pathways for cluster 4 proteins, which dramatically decreased in abundance after illumination. Though we did not find significantly enriched pathways containing photoreceptors, we also followed with interest the changes in the abundance of photoreceptors during the de-etiolation process. The abundances of PHYA, PHYB, PHYC and CRY2 were sharply downregulated after 12 hours of light treatment (Supplementary Table 1). This is consistent with the previous finding that photoreceptors are activated by light-induced phosphorylation, which eventually initiates their ubiquitination and degradation [17, 39, 40].

### Characterization of phosphorylated peptides

The number of phosphorylation sites per phosphorylated peptide and protein varied greatly. We found 8,746 phosphorylation sites in 9,528 phosphopeptides that matched 3,110 phosphoproteins (**Figure 5A**), and 7,118 (74.8%) peptides contained only one phosphorylation site (Figure 5B, Supplementary Table 2). Among the phosphorylated proteins, 1,057 (34%) contained more than three phosphorylation sites and 37 contained more than 16 phosphorylation sites (Figure 5C). For instance, the splicing factor PWI (GRMZM2G057646_P03) and cyclophilin (GRMZM2G006107_P02) were hyperphosphorylated, containing 48 and 40 phosphorylation sites, respectively (Supplementary Table 2). The most abundant phosphorylation site was S (7,639 or 77.78% of phosphosites), followed by T (1,067 or 12.20%) and Y (40 or 0.46%) (Figure 5A). This suggests that S is the chief site modified by phosphorylation in maize leaves.

**Figure 5.**
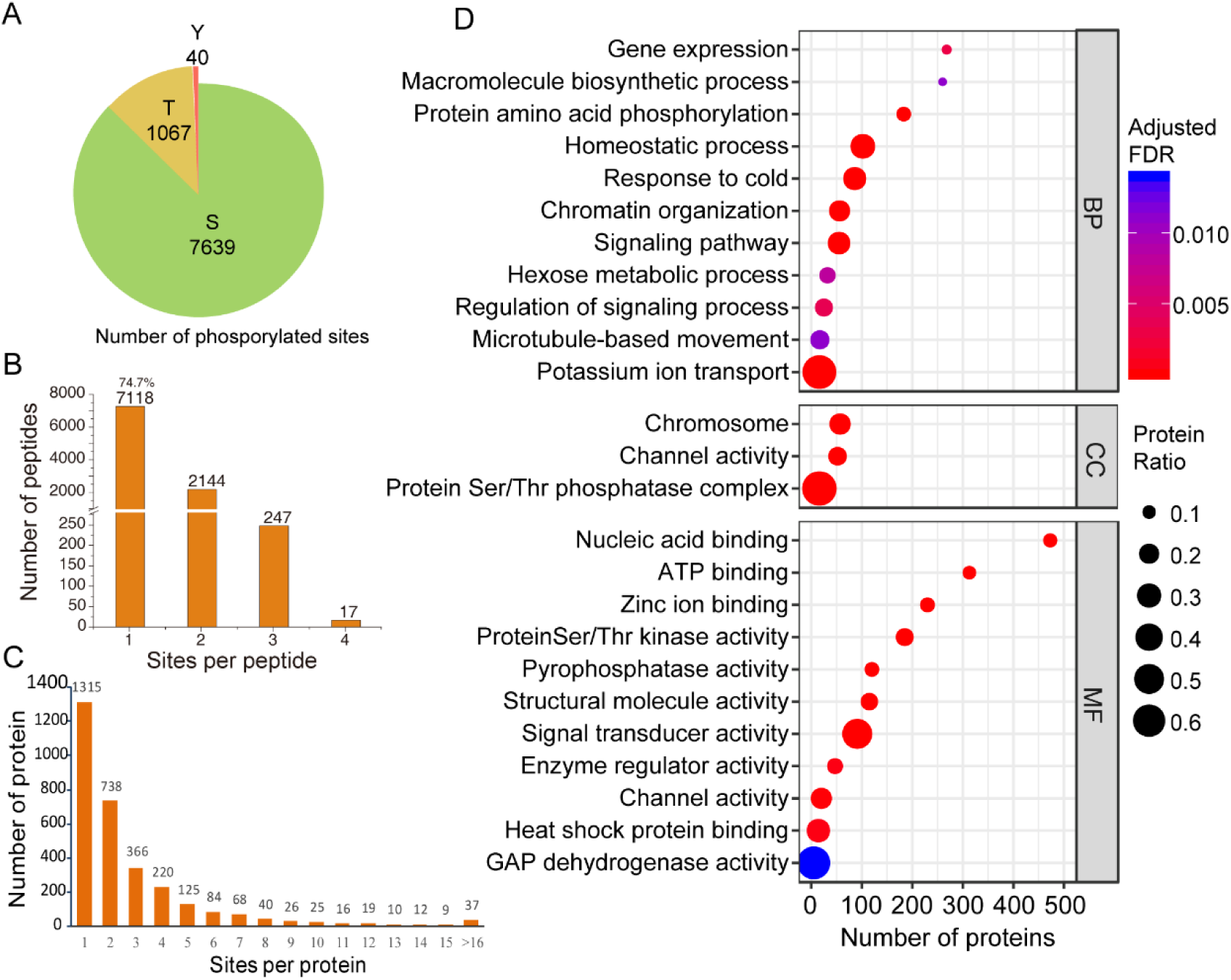
Overview of the phosphoproteome of de-etiolated maize seedling leaves. **A**. Pie chart showing the number of pS, pT, and pY phosphosites identified in the phosphoproteome. S accounts for most (87.33%) of the phosphosites. **B**. Distribution of phosphorylation sites per peptide; 74.7% of peptides with confidently assigned phosphosites were phosphorylated on a single site. **C**. Distribution of phosphoproteins. Over half of the phosphoproteins were phosphorylated on multiple sites. **D**. GO enrichment analysis of all phosphoproteins identified in this study. Based on GO slim terms all phosphoproteins were assigned to biological process (BP), cellular component (CC) and molecular function (MF) GO categories. Terms that were significantly enriched in DEPs (adjusted FDR≤ 0.05) are shown. The protein ratio is the ratio of the number of the phosphoproteins annotated to a certain term (adjusted FDR≤ 0.05) to the total number of proteins in the B73 maize genome assigned to that term. The horizontal axis indicates the total number of phosphoproteins annotated to each GO term. GAPDH: glyceraldehyde-3-phosphate dehydrogenase.

In the interest of revealing the pathways regulated by phosphorylation during the de-etiolation of etiolated maize seedlings, GO enrichment analysis of all 3,110 phosphoproteins was performed (Figure 5D). Eleven biological processes were highly enriched in phosphorylated proteins expressed during the de-etiolation process, such as protein amino acid phosphorylation, signaling pathway, and regulation of signaling process. The highest protein ratio (the number of phosphoproteins annotated to a certain GO term to the total number of proteins) of phosphoproteins was observed for the potassium ion transport term. This high ratio, 0.67, is because of the relatively low number of proteins (24) assigned to the potassium ion transport term. The ratios of phosphoproteins in the homeostatic process, response to cold, chromatin organization, and signaling pathway categories (0.31, 0.27, 0.22 and 0.25, respectively) were also relatively high. This indicates that phosphorylation modification may play a crucial role in the regulation of these pathways during the de-etiolation process.

To investigate which proteins bring about changes in phosphorylation during maize leaf de-etiolation, we screened our identified proteins for kinases and phosphatases (Supplementary Table 5). We quantified 816 protein kinases and 175 phosphatases, of which 234 kinases and 25 phosphatases were phosphorylated. A total of 543 kinases could be classified into 37 groups according to protein kinase classification system described by Lehti-Shiu and Shiu [41], and 40 phosphatases were classified into five families according to the ProFITS classification (http://bioinfo.cau.edu.cn/ProFITS/index.php), ITAK (http://bioinfo.bti.cornell.edu/cgi-bin/itak/index.cgi), and the P3DB database. These protein kinases and phosphatases include two plant-specific kinases, STN7/STT7 and STN8, and one phosphatase, TAP38/PPH1, that were previously shown to be involved in phosphorylation/dephosphorylation cycles in thylakoids associated with changes in light and diverse other environmental parameters [42]. Other kinases included those in the plant-specific TKL, STE, CMGC, and CK1 groups. We also identified calcium-dependent protein kinases (CDPK, 31), cyclin-dependent kinases (CDK, 23), mitogen-activated protein kinases (MAPK, 20), and MAPK cascade kinases (STE, 27), which play vital roles in transduction pathways (**Figure 6**, Supplementary Table 6).

**Figure 6.**
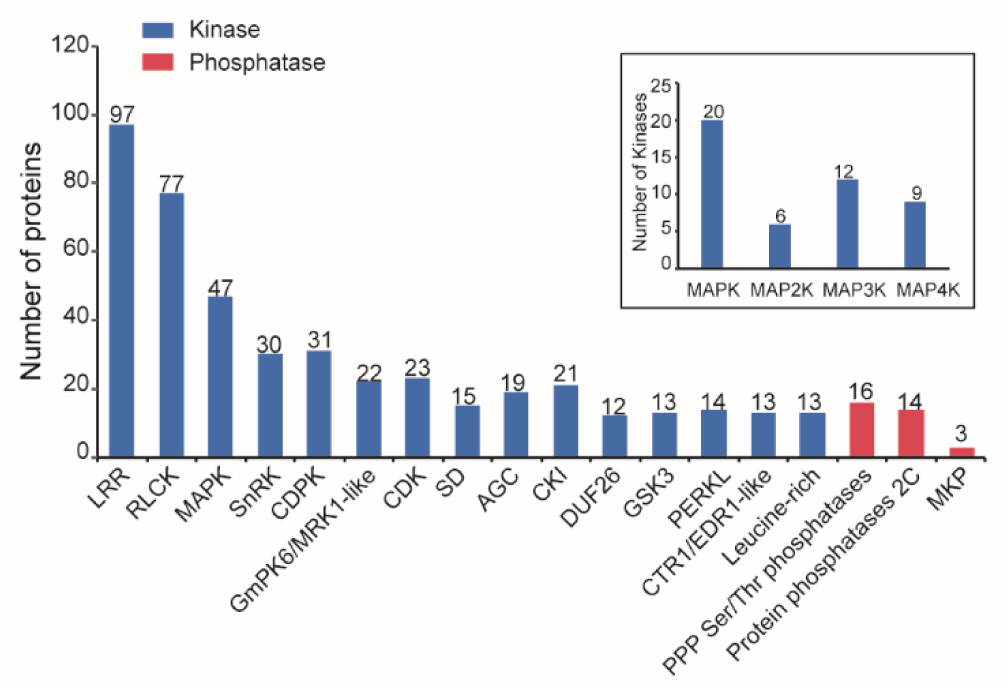
Classification of the identified kinases and phosphatases The kinases and phosphatases identified in this study were classified according to ProFITS (http://bioinfo.cau.edu.cn/ProFITS/index.php), ITAK (http://bioinfo.bti.cornell.edu/cgi-bin/itak/index.cgi), and the P3DB database. In total, 816 kinases and 175 phosphatases were identified. The kinase families including more than 10 members and the chief phosphatase families are shown. The inset shows the four MAPK families and the number of members of each family identified in this study. More information about the kinases and phosphatases is shown in Data S6

Predicted kinase-motif interactions and protein quantification/phosphorylation analysis can provide the basis for identifying possible substrates of different kinases. To identify the pathways that are potentially regulated by protein phosphorylation, we also identified phosphorylation motifs and the kinases that potentially phosphorylate these sites. Kinases that phosphorylate phospho-motifs where S or T is the central amino acid were classified into three major subgroups, namely proline-directed (pro-directed), basophilic (basic), and acidophilic (acidic), based on the types of substrate sequences preferred [43], and also into other families that we collectively refer to here as “other” (Supplementary Table 7). The pS- and pT-containing sites (99.6%) were also classified as pro-directed (33.82%), acidic (32.25%), basic (12.06%), and other (21.47%) (**Figure 7**). In this study, only 40 phosphorylated tyrosine peptides were identified, accounting for 0.46% of all phosphopeptides (Figure 7).

**Figure 7.**
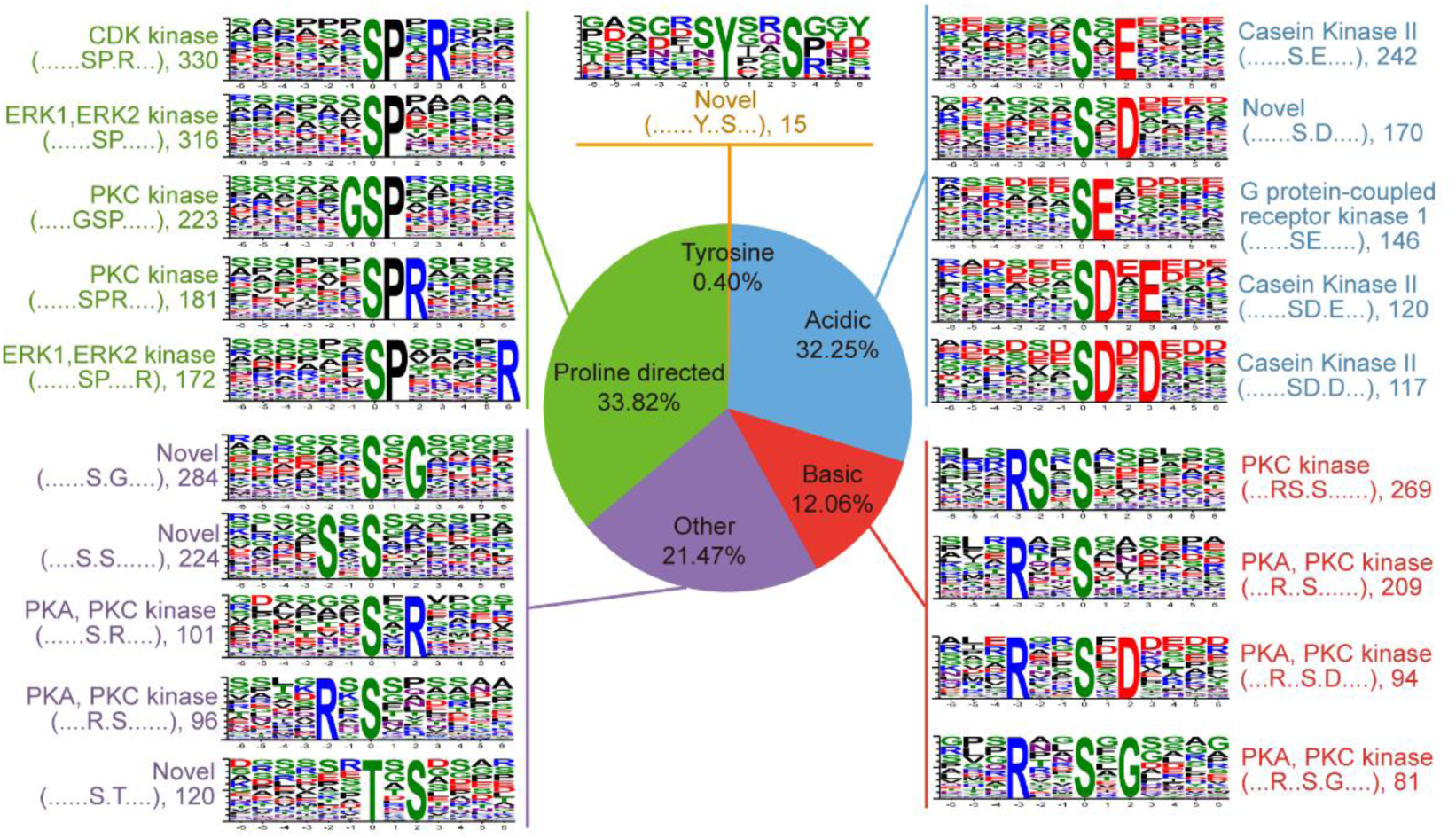
Motif classification and tabulation of known motifs Single phosphorylation motifs were identified using the Motif-X algorithm [78] and overrepresented motifs were extracted. The background was the *Zea mays*.AGP v3.28 protein database. The width was set to 13, the significance was set to 1×10^-6^, and the occurrence was set to 20 for serine and threonine motifs and to 15 for tyrosine motifs. The number shown after the type of motif indicates the number of times that the motif was identified. Sequence logos were generated with Weblogo (http://weblogo.berkeley.edu). Motifs were matched to known kinases using the Phosida motif matcher (http://phosida.de/) and the phosphomotif finder in the HPRD database (http://www.hprd.org/phosphomotif_finder) [44].

Using a method described previously [44], we identified phosphorylation motifs centered on S, T or Y residues that were overrepresented in phosphopeptides using motif-X (http://motif-x.med.harvard.edu) with the *Zea mays* AGPv3.28 protein database as the background. Using stringent criteria for S and T and looser criteria for Y, we identified 64 pS motifs, 13 pT motifs and 1 pY motif (Figure 7, Supplementary Table 7). The kinases that potentially phosphorylate these motifs were identified as described in the Materials and Methods. Casein kinase II substrates (sxE, sDxE, and sDxD) and basic motifs (RSxs, Rxxs, RxxsxD, and RxxsxG), which are phosphorylated by PKA and PKC kinases, were identified in our phosphopeptide dataset. Pro-directed substrates, which are mainly recognized by CDKs and MAPKs, were predominantly found among the phosphopeptides containing S phosphorylation sites (Figure 7, Supplementary Table 7). There were 330 phosphopeptides containing the pro-directed motif sPxR, which is recognized by CDK kinases. Pro-directed motifs sP and sPxxxxR, which are putatively phosphorylated by MAPKs, were found in 316 and 172 phosphopeptides, respectively. RBR1 (GRMZM2G0030343) contains sPxR, which can be phosphorylated by CDKA; 1 during the G1 to S phase transition [45]. The dataset of motifs and their corresponding kinases that we have generated can be used to identify new phosphorylation pathways, which will lead to a better understanding of the effect of phosphorylation on maize development.

### Phosphoproteome dynamics in maize leaves

The phosphorylation modification of proteins may be highly complex. Some phosphoproteins have multiple phosphosites, and phosphorylation may occur at different phosphosites under different conditions. There may also be differences in the change in NPL of peptides during de-etiolation. Here we describe the change in HY5 phosphorylation as an example. We identified three isoforms of HY5 (GRMZM2G039828_P01, GRMZM2G137046_P01 and GRMZM2G171912_P01) in maize leaves, and three pS sites were conserved in all three isoforms (Supplementary Figure 2). Six phosphopeptides of GRMZM2G039828_P01 were identified, which contained a total of four phosphosites. We identified two phosphorylated forms of the peptide “TSTTSSLPSSSER”. One was phosphorylated at the fifth S and the other was phosphorylated at the ninth S, and the changes in NPL were different for each peptide. Moreover, two other peptides containing the same pS or pT plus pS sites, which were possibly derived from different isoforms, showed very different changes in NPL.

In order to reveal what types of proteins are regulated by phosphorylation in etiolated seedlings exposed to light, we screened for significantly changed phosphopeptides (SCPPs) using stringent cutoff criteria (see Materials and Methods). In brief, the phosphopeptides with a fold change in NPL ≥1.5 or ≤0.67 were considered significantly changed. The proteins matching these peptides were considered proteins with significantly changed phosphorylation (PSCPs). We identified 1,475 PSCPs matching 826 proteins encoded by 823 genes. In fact, the number of PSCPs is likely much higher because many phosphorylated peptides were filtered out because the proteins they corresponded to were not quantified.

Firstly, we performed hierarchical clustering analyses of SCPPs using the average fold change in intensity ratios that were normalized by protein abundance (**Figure 8A**). 80% (1167) of the phosphopeptides, in which phosphorylation was immediately belonging to clusters 4, 5, and 6 increased after illumination. We also performed GO significantly enriched in nine biological pathways and two cellular components as PGKH, and iqd2. We next analyzed GO categories enriched in the phosphoproteins in each cluster 1 h and enrichment was observed for three BP categories (G-protein coupled receptor kinase activity, ATP binding, and structural molecule activity. These results suggest that the phosphorylation modification of proteins involved in the response to signaling pathway and kinase activities is affected by light and that changes in phosphorylation in response to light may regulate the de-etiolation process.

**Figure 8.**
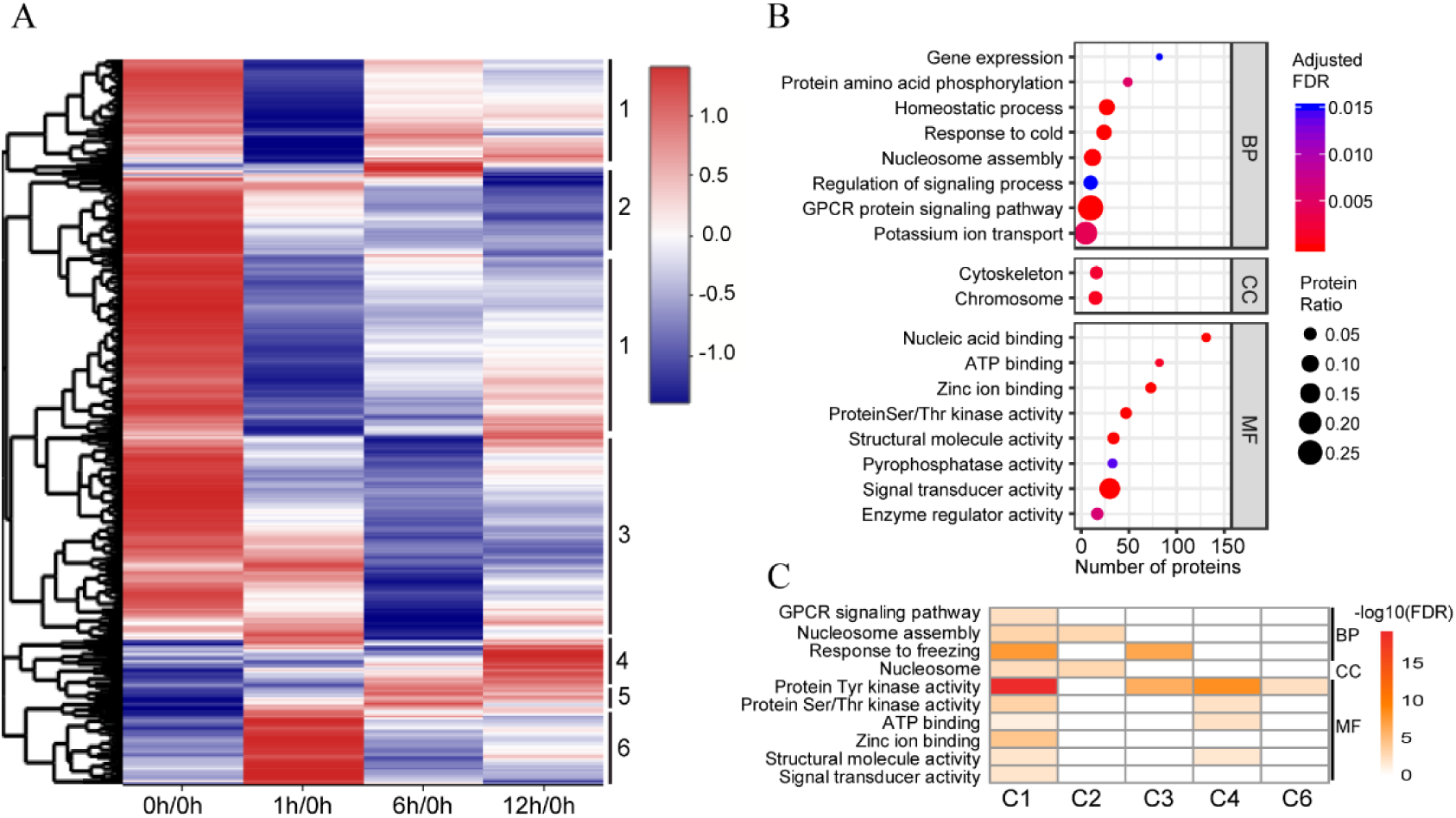
Phosphoproteome dynamics. **A**. Heat map of significantly changed phosphopeptides (SCPP). SCPPs were assigned to six clusters, which are shown on the right. The blue color represents a low level of phosphorylation while the red color represents a high level of phosphorylation. Vertical black lines on the right indicate the six clusters defined based on change pattern of NPLs. **B**. GO enrichment analysis of the proteins with significantly changed phosphorylation (PSCP) identified in this study. Based on GO slim terms, all PSCPs were assigned to biological process (BP), cellular component (CC) and molecular function (MF) GO categories. Terms that were significantly enriched in PSCPs (adjusted FDR≤ 0.05) are shown. The protein ratio is the ratio of the number of PSCPs annotated to a certain term (adjusted FDR≤ 0.05) to the total number of proteins in the B73 maize genome assigned to that term. The horizontal axis indicates the total number of PSCPs annotated to each GO term. **C**. GO enrichment analysis of the phosphoproteins in each cluster in A. The phosphoproteins were clustered into BP, CC, and MF GO categories. The color of each box indicates the –log10 (FDR) value. Yellow to red, significant enrichment; white, not significant. C1-C6: cluster 1-cluster 6; GPCR: G-protein coupled receptor.

### Transcription Factor dynamics

TFs play pivotal roles in the regulation of plant growth and development a but have traditionally been difficult to detect in proteomic analyses because of their low abundance [46, 47]. However, in de-etiolated maize leaves, we identified 724 (28.8%) proteins from 54 TF families comprising 2,516 annotated TFs listed in PlantTFDB 3.0 (Supplementary Table 8). The abundance of 37 (7.4%) out of 498 quantified TFs significantly changed during de-etiolation (Supplemental Table 8). Of these TFs, 29 were involved in the MapMan Bin “regulation of transcription”. Strikingly, the abundance of nine TFs significantly changed after 1 h of illumination (Supplementary Figure 4). Consistent with the role of the CONSTANS-like (CoL) protein CoL3 as a positive regulator of photomorphogenesis in *Arabidopsis* [48], we observed that the abundance of CoL4 and CoL5 drastically increased. We also observed dramatic changes in the abundances of one BES1 and two GATA proteins, which are considered to play central roles in the brassinosteroid signaling pathway and light signaling [49, 50].

Among all identified TFs, 209 were phosphorylated. Of these phosphorylated TFs, 77 were only identified through the enrichment of phosphorylated peptides, probably due to their low abundance (Figure 2B). The phosphorylation levels of 72 phosphopeptides matching 48 proteins belonging to 21 TF families significantly changed during de-etiolation (Supplementary Figure 4). Interestingly, the phosphorylation levels of two CoL proteins (CoL11 and CoL16) changed during de-etiolation. Therefore, our data suggest that these TFs with significant changes in phosphorylation level might function at higher levels in the hierarchy of gene transcriptional regulation during de-etiolation.

### Phosphorylation plays an essential role in the regulation of light signalling

Seventy-four proteins involved in various light signaling pathways were quantified. Only 13 of these proteins drastically decreased in abundance during de-etiolation. The NPLs of 25 phosphosites in 10 proteins drastically changed (Supplementary Table 9), indicating that phosphorylation of these sites may play an important role in regulating light signaling pathways.

PHYA is the major photoreceptor under far-red light in *Arabidopsis*, nevertheless PHYB plays a primary role under red or white light with the aid of PHYA, PHYC, and PHYD [51]. Under red light, rice PHYA and PHYB make equal contributions to seedling photomorphogenesis, and both rice PHYA and PHYC are included in far-red light response [52]. The maize genome has six genes encoding PHY proteins, including two PHYAs (GRMZM2G157727 and GRMZM2G181028), two PHYBs (GRMZM2G092174 and GRMZM2G124532), and two PHYCs (GRMZM2G057935 and GRMZM2G129889). Here we observed that two PHYAs, one PHYB (GRMZM2G092174), and one PHYC (GRMZM2G057935) significantly decreased in abundance during photomorphogenesis. This suggests that these four photoreceptors are likely involved in repressing photomorphogenesis in etiolated seedlings and that the de-etiolation mechanism regulated by PHY genes is highly conserved among monocotyledonous plants. It is noteworthy that although the abundance of the PHYB protein GRMZM2G124532 only decreased slightly in response to light, the NPLs of Thr-8, Ser-12, Ser-49 and Ser-76 in this protein drastically increased. In *Arabidopsis*, phosphorylation on Ser-86 plays an important role in modulating PHY-controlled signaling by accelerating the inactivation of PHYB [53]. Ser-76 in PHYB (GRMZM2G124532) and Ser-86 in AtPHYB are located in the same domain (Supplementary Figure 5); however, further experiments are needed to confirm whether they have similar functions. Nevertheless, these PTMs in PHYB may play important roles in modulating red or far-red light signaling pathways in maize seedlings just as in *Arabidopsis*.

CRY1 and CRY2 are responsible for photomorphogenesis under blue and UVA light[54], and autophosphorylation is important for their functions [55]. Here we quantified four out of the five maize CRY1 proteins (GRMZM2G024739, GRMZM2G049549, GRMZM2G104262, and GRMZM2G462690) and the single CRY2 protein (GRMZM2G172152). Only CRY2 drastically decreased in abundance, and at the same time the NPLs of Ser-480 and Ser-483 in CRY2 drastically decreased (Supplemental Table 8). In *Arabidopsis*, phosphorylation on three serine residues (Ser-588, Ser-599, and Ser-605) in the CCE (CRY C-terminal Extension) domain of CRY2 determines its photosensitivity [56]. Amino acid sequence alignments between the *Arabidopsis* and maize CRY2 proteins revealed that Ser-480 of the maize CRY2 proteins is located in the CCE domain and corresponds to Ser-599 of *Arabidopsis* CRY2 (Supplementary Figure 6). This suggests that the regulatory mechanism controlling CRY2 photosensitivity is likely conserved among various plant species.

Non-phototropic hypocotyl 3 (NPH3) is a member of a large family of highly conserved plant-specific proteins that interact with phototropins [57]. Previous studies showed that NPH3 is phosphorylated in dark-grown seedlings; its dephosphorylation is stimulated by blue light and appears to be correlated with phototrophism [58, 59]. In *Arabidopsis*, three phosphosites on the NPH3 (S212, S222 and S236) were identified by immunoblotting analysis, which were phosphorylated under dark conditions [60]. Here we identified 11 members of the NPH3 family in maize, and the abundance of one of them was significantly downregulated in response to light. Moreover, eight of these NPH3 proteins were found to be phosphorylated, and the NPLs of five phosphopeptides decreased during de-etiolation. For example, phosphorylation was identified on 16 phosphosites in one NPH3 protein (GRMZM2G413113), and the NPLs of the peptides, QSPSQNQpSPKpTPSR and WLPDVAPPTpSSSASGR, were the lowest at 6 h. The drastic reduction in the NPLs of most phosphosites during seedling de-etiolation (Supplementary Table 9) is in agreement with the deductions of previous studies [59, 60].

### Phosphorylation plays an important role in regulating proteins involved in photosynthesis light reactions

During de-etiolation, proteins involved in photosynthetic pathways had the most dramatic increases in abundance. In previous studies, 13 of 52 significantly changed proteins in rice and 31 of 73 in maize were related to the photosynthetic pathway [24, 25]. Here, we identified 158 proteins involved in photosynthesis light reactions, and 154 of them were quantified (**Figure 9**, Supplementary Table 10). Of these proteins, 84 (54.5%), including 30 PSI and 37 PSII subunits, significantly increased in abundance during de-etiolation. For example, CP29 abundance was 1.99-, 6.76-, and 9.41-fold higher in the 1 h, 6 h, and 12 h samples, respectively, than in the 0 h sample.

**Figure 9.**
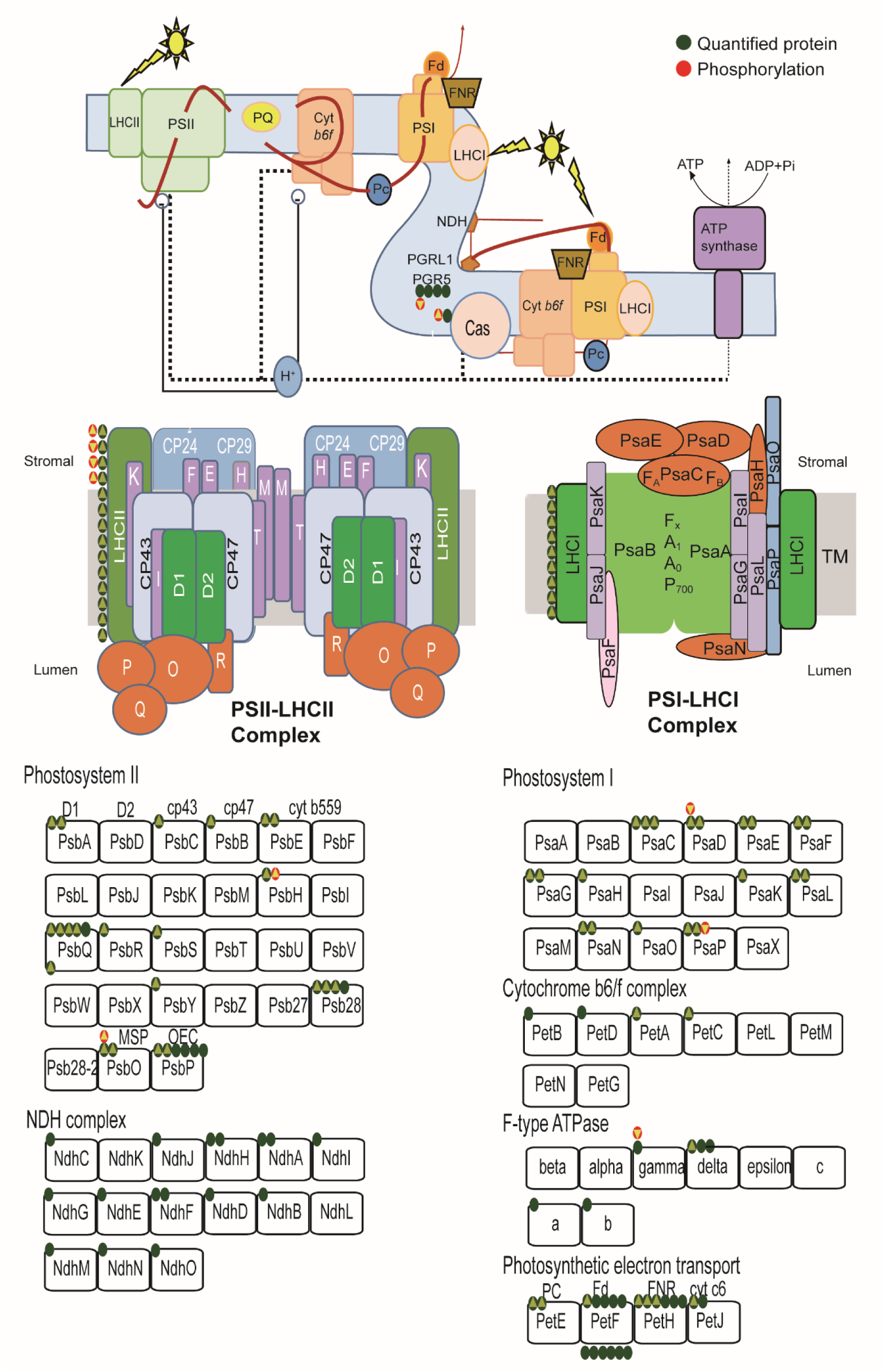
Phosphorylation of proteins involved in photosynthetic light reaction pathways changes significantly during de-etiolation This figure was modified from figures in Eberhard et al. (2008) and Nickelsen and Rengstl (2013) and the KEGG photosynthesis pathway (https://www.kegg.jp/kegg-bin/) [79, 80]. In the top of the figure, the complexes involved in the photosynthesis electron transfer chain are shown. A model for the assembly of PSII and PSI complexes is shown in the middle of the figure. The components contained in each complex are listed at the bottom of the figure. The green and red circles indicate quantified and phosphorylated proteins, respectively, that function in the light reactions of photosynthesis. The number of circles for each component represents the number of homologs identified in this study. A green circle with an upward triangle indicates that the abundance of the protein was significantly upregulated during de-etiolation; a red circle with an upward or downward triangle indicates that the NPL of the protein was significantly upregulated or downregulated, respectively, during de-etiolation.

Eleven proteins involved in photosynthesis light reactions were found to be phosphorylated, and the NPLs of nine proteins, namely four PSII subunits (PSbO, PSbH, CP29, and LHCB1.5), two PSI subunits (PSaD-1 and PSAP), an ATP synthase subunit (δ subunit), a protein involved in cyclic electron flow (CEF) (PGRL1), and a calcium sensing receptor (CAS), drastically changed (Supplemental Table 9). In particular, the NPL of Thr377 in CAS increased 2.8-fold at 1 h, 13.7-fold at 6 h, and 20.0-fold at 12 h compared with the 0 h sample (Supplemental Table 9). Sequence alignment indicated that Thr377 of maize CAS corresponds to Thr380 of *Arabidopsis* CAS, which is one of the target sites of the thylakoid protein kinase STN8 (Supplementary Figure 7). In *Arabidopsis*, CAS is essential for regulating the transcription of photosynthetic electron transport-related genes, the formation of the photosynthetic electron transport (PET) system, and water use efficiency [61]. Therefore, the drastically increased NPL of Thr377 in CAS might be tightly related to the formation or the regulation of the PET system.

### Phosphorylation plays a pivotal role in regulating the activities of enzymes involved in carbon assimilation

As a classical C4 plant, both the Calvin cycle and the C4 carbon assimilation pathway are active in maize leaves. Here, we identified 85 proteins involved in carbon assimilation, including 43 enzymes involved in the Calvin cycle and 42 C4 pathway enzymes (Supplemental Table 11). Of these proteins, 33 significantly increased in abundance during de-etiolation. Moreover, 17 proteins related to the Calvin cycle and the C4 cycle were phosphorylated, and the NPLs of 31 phosphopeptides corresponding to 12 of these proteins changed; these changes in NPL were significant for 21 of the peptides. These phosphoproteins with significant changes in NPL included key enzymes in the Calvin cycle, such as the Rubisco small subunits, phosphoglycerate kinase, and glyceraldehyde-3-phosphate dehydrogenase, and key enzymes in the C4 cycle, such as PPDK, PEPC, and PEPCK. For these proteins, the increase in NPL occurred in parallel with an increase in abundance (**Figure 10**, Supplemental Table 11). This suggests that phosphorylation of these proteins is likely related to the regulation of their enzymatic activities.

**Figure 10.**
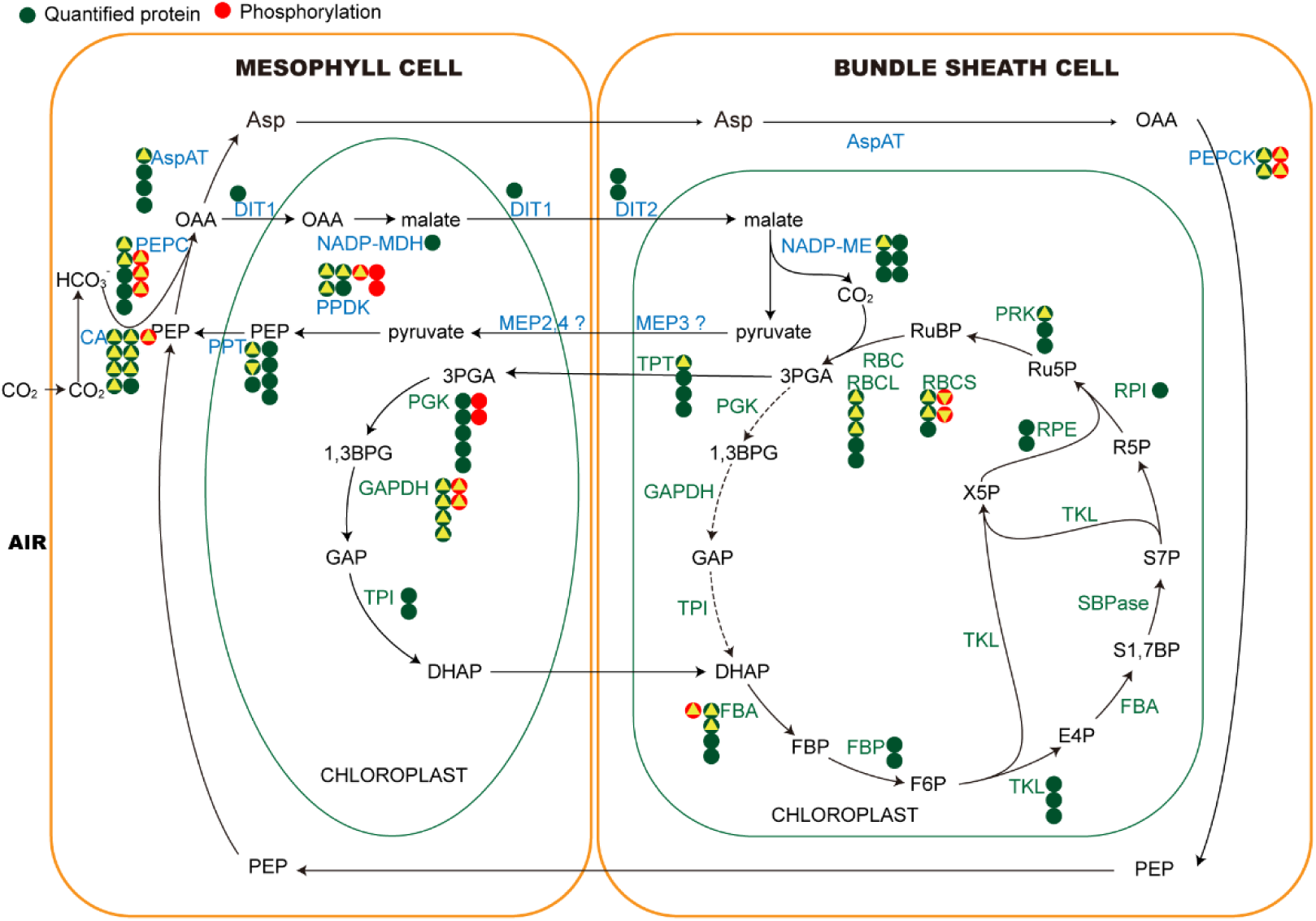
Phosphorylation of proteins involved in the C4 cycle of photosynthesis changes significantly during de-etiolation This figure was modified from figures in Majeran et al. (2008) and Wingler et al. (1999) [28, 65]. Proteins written in blue font belong to the C4 pathway, and proteins written in green font belong to the Calvin cycle. The green and red circles indicate quantified and phosphorylated proteins, respectively, that function in the light reactions of photosynthesis. The number of circles for each component indicates the number of homologs identified in this study. A green circle with an upward triangle indicates that the abundance of the protein was significantly upregulated during de-etiolation; a red circle with an upward or downward triangle indicates that the NPL of the protein was significantly upregulated or downregulated, respectively, during de-etiolation. CA: Carbonic anhydrase; PEPC: Phosphoenolpyruvate carboxylase; AspAT: Aspartate aminotransferase; DIT: Dicarboxylate transporter; NADP-MDH: Malate dehydrogenase [NADP]; NADP-ME: NADP-dependent malic enzyme; MEP: Envelope protein; PPDK: Pyruvate, phosphate dikinase; PPT: Phosphoenolpyruvate (PEP)/phosphate translocator; RBCL: large subunit of Rubisco; RBCS: small subunit of Rubisco; PGK: Phosphoglycerate kinase; GAPDH: Glyceraldehyde-3-phosphate dehydrogenase; TPI: Triosephosphate isomerase; FBA: Fructose-bisphosphate aldolase; FBP: Fructose-1,6-bisphosphatase; TKL: Transketolase; SBPase: Seduheptulose bisphosphatase; RPI: Ribose-5-phosphate isomerase; RPE: Ribulose-phosphate 3-epimerase; PRK: Phosphoribulokinase; PEPCK: Phosphoenolpyruvate carboxykinase.

PEPC is one of the most important C4 photosynthesis enzymes in maize and catalyzes the carboxylation of phosphoenolpyruvate (PEP) to yield oxaloacetate and inorganic phosphate [62]. The NPLs of two sites in three PEPC proteins (GRMZM2G069542_P01, GRMZM2G083841_P01 and GRMZM5G845611_P01) were drastically upregulated in response to light at 1 h compared with those at 0 h, and the NPL of Ser15 in GRMZM2G083841_P01 increased 11-fold (Supplementary Table 11). After 6 h of illumination, the NPLs of these three PEPC proteins decreased to a level similar to that of the 0 h sample, which suggests that phosphorylation rapidly adjusts the enzyme activities of PEPC proteins in response to light.

PPDK is one of the key enzymes involved in C4 photosynthesis and plays an important role in regenerating PEP. Moreover, the activity of PPDK may limit the rate of CO2 assimilation in the C4 cycle [63]. We previously reported that light intensity regulates PPDK activity by modulating the reversible phosphorylation of Thr-527 by the PPDK regulatory protein (PDRP) [64]. The phosphorylation of Thr-527 inhibits the enzymatic activity of PPDK. Here, the NPL of Thr-527 in PPDK (Thr-463 in GRMZM2G097457_P02 and Thr-462 in GRMZM2G306345_P03) was significantly downregulated after 6 h of illumination. This implies that the activity of PPDK was enhanced and the rate of the carbon reaction increased late during de-etiolation. This may be due to the rapid progression of the light reaction after the establishment of the photosynthetic machinery.

In maize leaves, PEPCK is mainly located in bundle sheath cells and participates in CO_2_ concentration by catalyzing the conversion of oxaloacetate to PEP, releasing one CO_2_ [65]. The NPLs of eight different sites (Ser47, Thr50, Thr51, Ser55, Thr58, Thr59, Ser67, and Thr120) in PEPCK drastically changed after 6 h of illumination. The NPLs of Ser67 and Thr120 were sharply downregulated, while those of the other six sites were significantly upregulated. One peptide containing four phosphorylated sites (Ser47, Thr51, Thr58, and Thr59,) showed a 10-fold increase in NPL.

### Phosphorylation of plasma and plastid proteins plays an important role in metabolism and the flow of ions during the de-etiolation of etiolated seedlings

Transporters are located on both the plasma and plastid membranes, and the combined activity of all transporters regulates de-etiolation. During the de-etiolation of maize etiolated seedlings, sugar, auxin, ABA, cation and anion transporters are phosphorylated, and this modification affects their activities [23, 24, 30]. To analyze the change in transporter abundance and phosphorylation during de-etiolation, we screened the de-etiolated maize seedling proteomic and phosphoproteomic data set for sugar, hormone, ion (especially calcium transporters in the chloroplast), amino acid, and ammonium transporters. Of the 730 transporters identified 565 were quantified. Of the 47 quantified proteins with significant changes in abundance during de-etiolation (Supplementary Table 12), 34 were significantly upregulated. For example, the abundance of Cationic amino acid transporter 9 (CAT9, GRMZM2G139920) at 6 h was more than 4.8-fold higher than the abundance at 0 h (Supplementary Figure 8). Three out of six putative SWEET family proteins (GRMZM2G144581, GRMZM2G106462, and GRMZM2G111926) and one plastid glucose transporter (pGlcT1, GRMZM2G098011), which are intercellular and chloroplast sugar transporters, respectively [66, 67], were upregulated. In addition, 116 transporter proteins were found to be phosphorylated, and the NPLs of 49 sites in 26 proteins changed significantly.

Tonoplast monosaccharide transporter1/2 (TMT1/2) proteins are sugar transporters that are localized on the vacuolar membrane and probably load glucose and sucrose into the vacuole [68]. ERD6-like proteins are involved in the transport of sugars out of the vacuole during conditions such as wounding, pathogen attack, senescence, and Carbon/nitrogen (C/N)-starvation and play roles opposite to those of TMT1/2 [69, 70]. In *Arabidopsis*, four TMT2 phosphorylation sites are located in the central hydrophilic loop, which was previously reported to be more heavily phosphorylated after cold induction, resulting in enhanced TMT activity [71]. ERD6-like can also be phosphorylated at S residues in *Arabidopsis* [70]. In the present study, NPLs of four sites in TMT2 (Ser-276, −280, −286 and −320 in GRMZM2G083173) were downregulated, while the NPL of Ser-55 in the ERD6-like (GRMZM2G097768) protein was upregulated. In contrast, no change in protein abundance was observed in Schulze’s study or in our study. This may be because the regulation of transporter activity by phosphorylation may allow more rapid regulation of sugar transport than changing protein abundance.

Metal ions are critical cofactors for legion chloroplast proteins involved in photosynthesis (Ca^2+^, Mg^2+^, Mn^2+^ and Fe^2+^) and oxidative stress detoxification (Cu^2+^, Zn^2+^ and Fe ^2+^), and H^+^-coupled ATPases are important for chloroplast biogenesis. The activity of metal ion transporters controls the concentration of ions in specific locations.

For example, CCHA1 (GRMZM2G150295) is a chloroplast-localized Ca^2+^/H^+^ antiporter, which plays critical roles in containing chloroplast Ca^2+^ and pH homeostasis and the regulation of PSII in *Arabidopsis* [72], but the PTM of CCHA1 has not been studied so far. To understand whether the activity of these transporters is potentially regulated by phosphorylation, we screened for phosphorylation of these cation and proton pumps. We found that the NPLs of Thr7 and Ser8 in CCHA1, Ser298 in CAX interacting protein 4 (CXIP4, GRMZM2G048257), Ser82 in potassium transporter KUP12 (GRMZM2G036916) and Thr881 in autoinhibited H^+^-ATPase isoform 2 (AHA2, GRMZM2G019404) were significantly upregulated during de-etiolation. The phosphorylation modifications identified in the present study could provide guidance for studies of the regulation of the functions of these transporters by PTMs.

## Conclusion

The transition from etiolation to de-etiolation is a very complicated process, during which plants need to quickly respond to light signals and rapidly mobilize photomorphogenesis to complete the formation of the photosynthetic system and initiate photosynthetic reactions. In the present study, we have provided the most comprehensive dynamic analysis of protein abundance and phosphorylation in de-etiolated maize leaves to date.

Among the quantified proteins, the abundance of only 6%, including 37 TFs, significantly changed during the de-etiolation process; these proteins include nearly all those previously shown to change in abundance at similar developmental stages [23, 25, 73]. In contrast, the phosphorylation levels of 26.6% of the quantified proteins, especially those involved in gene expression, protein amino acid phosphorylation, and homeostatic process pathways, significantly changed. In addition, 25.2% out of all identified phosphorylated peptides contained more than two phosphosites; these peptides corresponded to 1,057 (34%) phosphoproteins containing three or more phosphosites, and 128 of them contained more than 10 phosphosites. These phosphosites may regulate different aspects of protein function by activating or inhibiting protein activity, which may in turn regulate the functions of these proteins in different pathways. Moreover, these effects may be enhanced by phosphorylation at multiple sites in the same protein.

Our data suggest that the regulation of PTM levels on proteins might be more efficient than the regulation of protein abundance for adapting to changing environments. Reversible PTMs allow plants to rapidly respond to internal and external cues. In addition, PTM is more economical in terms of energy use than transcriptional regulation, which involves several steps from the initiation of gene transcription to the formation of a mature protein; only a little energy (ATP or GTP) is needed to add or to remove a functional group (PTM) on a protein in order to change its physical and chemical properties. Therefore, the study of protein PTM is important to fully explore the mechanisms of plant adaptation to environmental changes.

## Materials and Methods

### Plant material and sample collection

The maize inbred line B73 was used in this study. The seedlings were planted and samples were collected as described previously [32]. Under the same conditions, two biological replicates were performed and the first seedling leaves from each replicate were rapidly sampled. All samples were frozen in liquid nitrogen and stored at −80 °C until further use.

### Protein extraction

Total proteins were extracted from maize seedling leaves using a 10% (w:v) trichloroacetic acid (TCA)/acetone solution as described previously [32]. The protein concentration of each sample was determined using the 2-D Quant kit (GE Healthcare). Protein samples were stored at −80 °C for further experiments.

### Sample preparation

Protein extraction of two sets of maize samples (0 h, 1 h, 6 h, and 12 h, ∼5 mg each) and trypsin digestion were performed as previously described [32].According to the manufacturer’s instructions, samples were labeled with iTRAQ 4plex reagent (ABSciex, MA, US) and then combined.

### Strong Cation Exchange (SCX) Chromatography

Total iTRAQ-labeled lysate was solubilized in buffer A (5 mM KH2PO4/25% acetonitrile, pH 3.0) and separated on a PolySULFOETHYL A column (4.6 mm ID x 100mm, 5 µm, 300 Å, Poly LC Inc, Columbia, MD) with flow rate of 1 ml/min using a linear gradient of 0% buffer B (5 mM KH2PO4/25% acetonitrile/400 mM KCL, pH 3.0) to 100% buffer B over 40 min. A Gilson system composed of 306 pumps, an 805 manometric module, an 811C dynamic mixer, and a UV/VIS-155 detector was used. The sample fractions were collected every minute and dried.

### Basic reverse phase HPLC

For general proteomics, the iTRAQ-labeled total lysate or the selected SCX fractions described above were solubilized and separated as previously described [74]. For phosphoproteomics, ammonium formate was switched to ammonium every 6th fraction from fractions 10-45 to generate a total of 6 pooled fractions. The pooled fractions were subsequently dried under vacuum and subjected to phospho-peptide enrichment using the IMAC method or TiO_2_.

### Phosphopeptide enrichment using IMAC and TiO_2_

The protocol for IMAC enrichment of phosphopeptides was adapted from Mertins et al. with modifications [75]. The procedure for phosphopeptide enrichment using TiO_2_ was adapted from Wilson-Grady et al. with modifications [76]. The enriched phosphopeptides were further desalted using an Empore 3M C18 (2215) StageTip^2^ prior to HPLC-MS/MS analysis.

### HPLC-MS/MS

A Dionex RSLC system interfaced with QExactive HF (ThermoFisher, San Jose, CA) was used to carried out HPLC-MS/MS primarily. Due to instrument availability, 2D-HPLC-MS/MS and phosphor-proteomic samples were analyzed using a Dionex RSLC system interfaced with a Velos LTQ Orbitrap ETD (ThermoFisher, San Jose, CA) as described previously [32]. To achieve the mass spectrometry data, a data-dependent acquisition procedure with a cyclic series of full scans was used and acquired in the Orbitrap with a resolution of 120,000 (QExactive HF) or 60,000 (VELSO LTQ Orbitrap ETD). MS/MS scans of the ten most intense ions were followed then and scanned out in the Orbitrap with a resolution of 30,000 (QExactive HF) or 15,000 (VELOS LTQ Orbitrap ETD) with low mass set at 110 amu.

### Data analysis

The HPLC-MS/MS data from each experiment were searched as described previously [32]. For proteins identified only in 2D-HPLC-MSMS, the average of the ratios from two biological replicates was used to represent the final fold change at each time point, while for proteins identified only in 3D-HPLC-MSMS or both in 2D and 3D, the 3D ratios were used. The proteins were considered significantly changed with a fold change ≥1.5 or ≤0.67.

### Database searching with the phosphoproteome data

The LC-MSMS data were searched in MUDPIT style against the *Zea mays* database using MASCOT (version 2.3 MatrixScience, UK) or MSAmanda (version 1.4.14.3866 for PD1.4.1.14) [77] in the Proteome Discoverer (version 1.4.1.14, Thermo Fisher Scientific, Bremen, Germany) environment. For both search engines, oxidation of methionine and phosphorylation on serine, threonine and tyrosine were set as variable modifications. MASCOT and MSAmanda results were combined for reporting. Percolator was used to validate PTMs. Only top hit peptides with FDR<0.01 (based on PEP score) were included in the final results. Phosphorylation sites were localized using PhosphoRS 3.1 (implemented in Proteome Discoverer 1.4.1.14). The ratio of phospho-peptides between the 0 h control and different time points were calculated using reporter ion intensities and further normalized to general normalization factors determined from the median of high confidence spectra from HPLC-MS/MS results obtained prior to enrichment (fractionation-phospho-peptide enrichment) or general proteomic data (phospho-peptide enrichment from the total lysate) as described in the general proteomic section of the Materials and Methods. For the phosphopeptides only quantified in one experiment, the ratios were used to represent the final fold change at each time point, while for the peptides quantified more than twice, the mean value of the ratios from all replicates was used as the final fold change. The phosphopeptides with a fold change ≥1.5 or ≤0.67 were considered significantly regulated.

### SDS-PAGE and Immunoblotting

Protein expression and phosphorylation were assessed using standard western blotting protocols described by Chen et al. [64]. Blots were probed with rabbit polyclonal anti-AtpB, anti-PsaD, anti-PsbC, anti-RA, anti-RbcL, anti-RbcS and anti-UGPase antibodies (Agrisera Antibodies, Vännäs, Sweden) and an anti-plant-actin rabbit polyclonal antibody (EasyBio, Beijing, China). The rabbit polyclonal anti-PPDK, anti-PEPCK and phosphosite-specific anti-PT527 PPDK antibodies were prepared by our laboratory.

### Motif and kinase-phosphatase analysis

Phosphopeptide sequences were extended to 13 aa with a central S, T or Y using the *Zea mays* database (V3.28) [78]. Pre-aligned peptides were submitted to the Motif-X algorithm (http://motif-x.med.harvard.edu/). Sites that were located at the N- or C-terminus and could not be extended to 13 aa were excluded. The significance was set to P<10^-6^, and the minimum number of motif occurrences was set to 20 for S and T and to 15 for Y. Motifs were classified as proline-directed, acidic, basic and other as described previously [44]. Sequence logos were generated with Weblogo (http://weblogo.berkeley.edu). All proteins identified in this study were screened for kinases and phosphatases using the GO accession number GO: 0016301 for kinases and GO: 0016791 for phosphatases. Kinases and phosphatases were classified according to the maize databases ProFITS (http://bioinfo.cau.edu.cn/ProFITS/index.php), ITAK (http://bioinfo.bti.cornell.edu/cgi-bin/itak/index.cgi), and P3DB (http://p3db.org/).

### Bioinformatics analyses

Protein sequences were obtained from EnsemblPlants (http://plants.ensembl.org/index.html). The functional annotations of protein were performed using the MapMan (*Zea Mays* genome release 1.1) and Ensemble Plant (AGPv3) databases.

Hierarchical clustering of proteins was done in R (version 3.4.3; https://www.r-project.org) using the heatmaply method from the heatmaply package (version 0.14.1). GO enrichment analysis of proteins and phosphoproteins was performed based on agriGO v2.0 (http://systemsbiology.cau.edu.cn/agriGOv2/index.php) with the B73 maize genome V3.3 as the background. Significantly enriched terms in the BP, CC and MF GO categories were plotted using ggplot from the ggplot2 package (version 2.2.1; http://ggplot2.tidyverse.org) in R (version 3.4.3; https://www.r-project.org).

### Sequence analysis

Sequences of HY5, CRY2 and CAS from *Arabidopsis* and maize were downloaded from the Phytozome database (https://phytozome.jgi.doe.gov/pz/portal), and sequence alignment was done using BioEdit software.

### Data availability

Accession number: these mass spectrometry proteomics and phosphoproteomics data have been submitted to the ProteomeXchange Consortium (http://proteomecentral.proteomexchange.org/cgi/GetDataset) with the dataset identifier PXD012897.

### Authors’ contributions

BCW, QC, ZFG and ZS conceived of this study and participated in its design. ZFG performed statistical analyses and wrote the manuscript. ZS prepared plant materials, extracted proteins and performed some of the statistical analyses. QC gave suggestions on the manuscript and performed manuscript revision. ZY collected the antibodies and did the western blotting analysis. XLG performed searches and analysis of network databases in the manuscript. TL processed proteomics data and composed part of the experimental procedures in the manuscript. HZ performed the MS experiments. All authors read and approved the final manuscript.

## Acknowledgments

This work was supported by the National Key Research and Development Program of China (Grant No. 2016YFD0101003) and Heilongjiang Provincial Outstanding Youth Science Foundation (JC2017008, Heterosis characteristics and regulation mechanism of nitrogen physiological utilization efficiency of maize under different nitrogen supply levels). We thank Dr. Ke Cao for providing guidance on the use of R language to generate images of the results of hierarchical clustering analyses and GO enrichment. We thank Lehti-Shiu Melissa (Lehti Life Science Editing) for offering service for proofreading the manuscript.

## Supplementary material

**Data S1 Proteome**

**Data S2 Phosphoproteome**

**Data S3 The proteins used for drawing Venn diagrams**

**Data S4 Total proteins**

**Data S5 Clusters of DEPs**

---

**Supplementary Figure 1.**
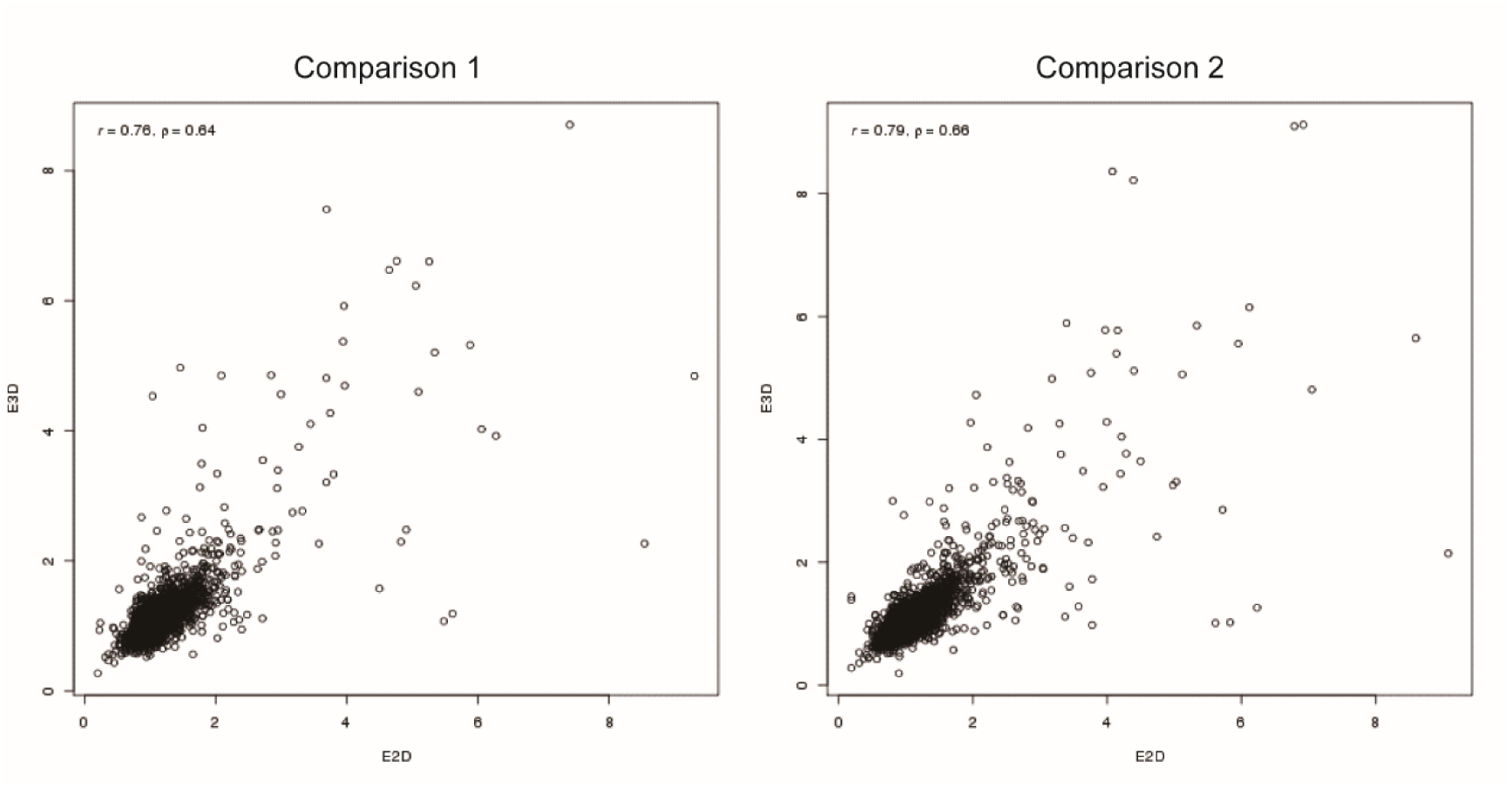
Comparisons between the 2D proteome and 3D proteome. The protein abundance data from the 3D proteome analysis were compared with the data from the first (Comparison 1) and second (Comparison 2) repeats of the 2D proteome analysis. The Y-axis is the ratio of 1 h/0 h, 6 h/0 h and 12 h/0 h identified in 3D proteome analysis and the X-axis is the ratio of 1 h/0 h, 6 h/0 h and 12 h/0 h identified in the 2D proteome analysis. The Pearson correlation coefficient (r) and the Spearman correlation coefficient (ρ) are shown in the upper-left corner of each plot.

**Supplementary Figure 2.**
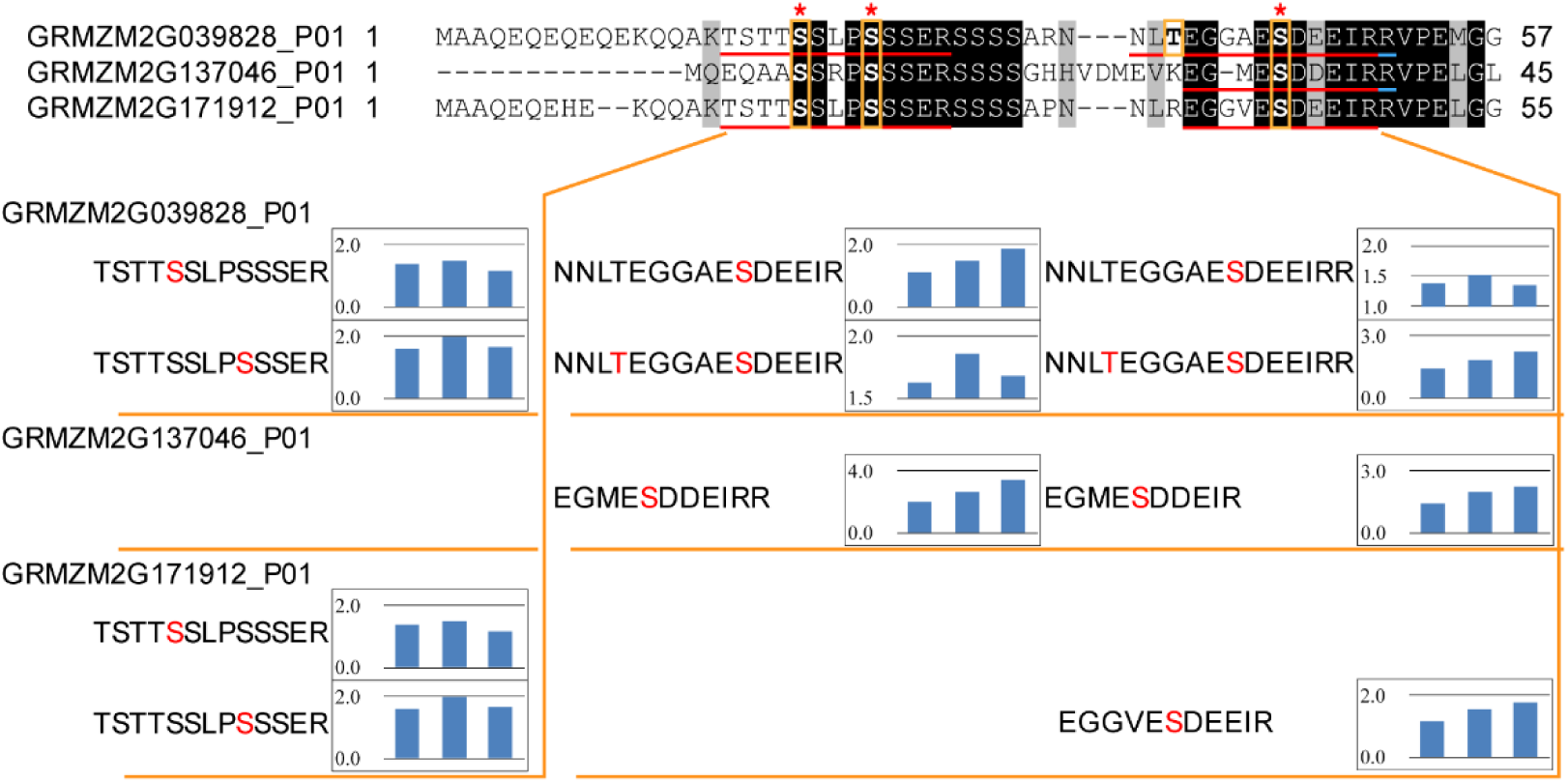
The phosphorylation of three HY5 isoforms in maize. Peptides corresponding to three HY5 proteins that contain phosphorylation sites identified in this study are shown at the top of the figure. The “S” and “T” residues highlighted by yellow boxes are the phosphorylation sites, and the S residues are conserved in all three peptides. The phosphorylated peptides identified by HPLC-MS/MS after enrichment using IMAC and TiO_2_ are shown below the sequence alignment, and the red S and T residues are the phosphorylation sites. The bar graphs to the right of each peptide show the changes in NPL after illumination for 1, 6 and 12 hours. The red asterisks indicate the main phosphorylation sites.

**Supplementary Figure 3.**
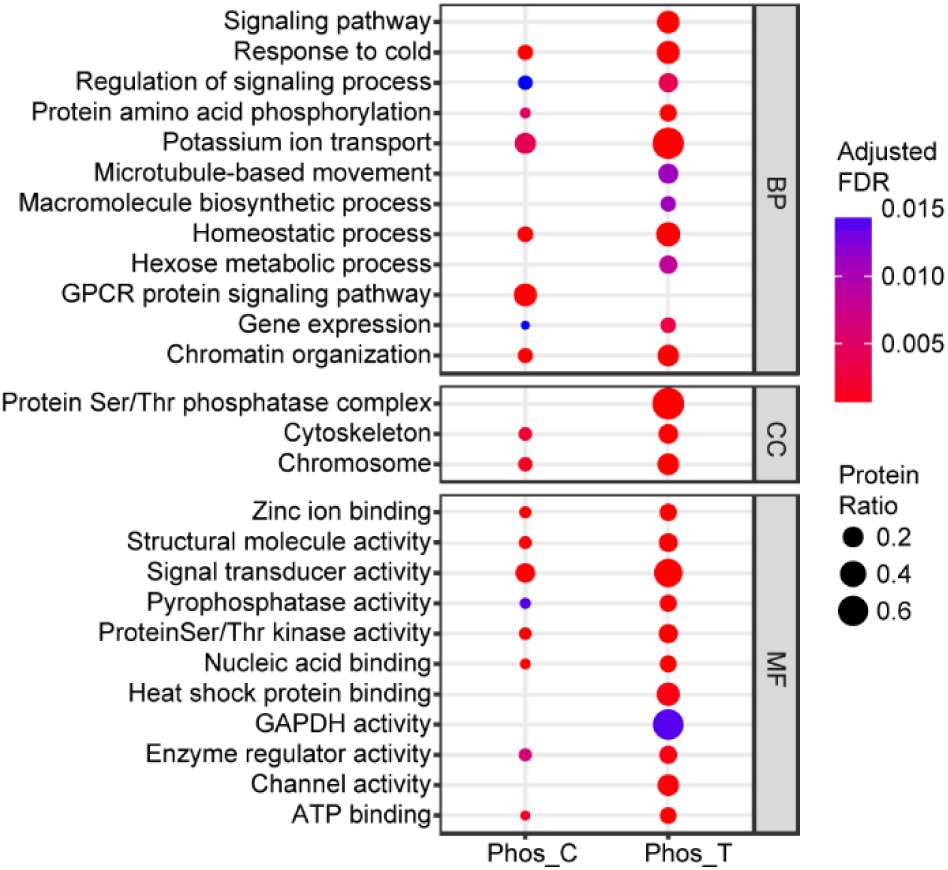
Enrichment analysis of all phosphoproteins and the phosphoproteins with significantly changed phosphorylation Based on GO slim terms, all phosphoproteins (Phos_T) and differentially phosphorylated phosphoproteins (Phos_C) were assigned to biological process (BP), cellular component (CC) and molecular function (MF) GO categories. Terms that were significantly enriched in phosphoproteins and differentially phosphorylated phosphoproteins (adjusted FDR≤ 0.05) are shown. The protein ratio is the ratio of the number of phosphoproteins or differentially phosphorylated phosphoproteins annotated to a certain term (adjusted FDR≤ 0.05) to the total number of proteins in the B73 maize genome assigned to that term. GPCR protein signaling pathway: G-protein coupled receptor protein signaling pathway; GAPDH: glyceraldehyde-3-phosphate dehydrogenase.

**Supplementary Figure 4.**
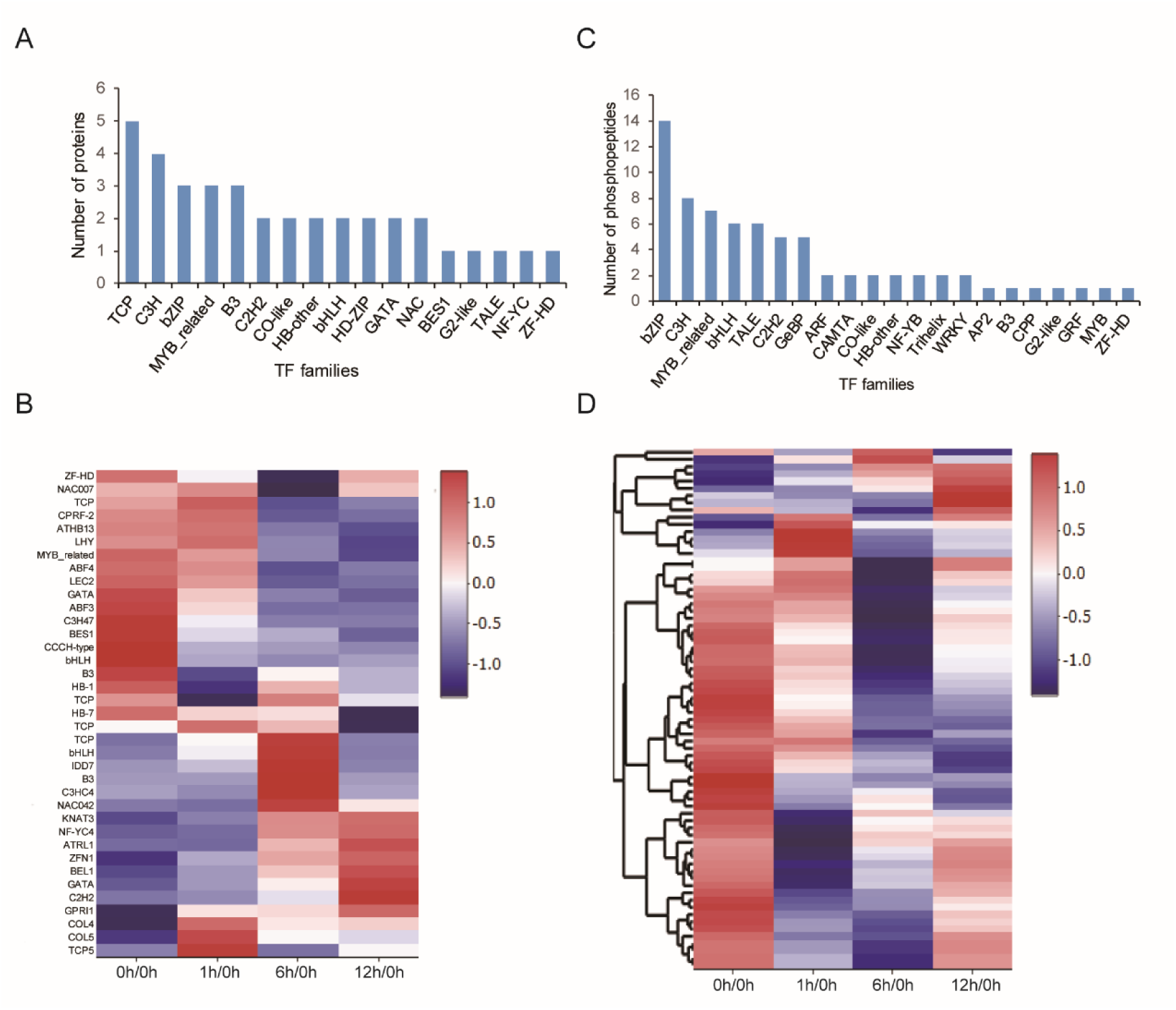
The dynamics of the TFs identified in the proteome and phosphoproteome **A**. The number of TFs in 17 families with significant changes in protein abundance are shown. **B**. Heat map of the TFs that significantly changed in abundance during de-etiolation. **C**. The number of TFs in 21 families with significant changes in normalized phosphorylation levels. **D**. Heat map of the TFs with significant changes in normalized phosphorylation levels during de-etiolation. The blue color represents low abundance while the red color represents high abundance.

**Supplementary Figure 5.**
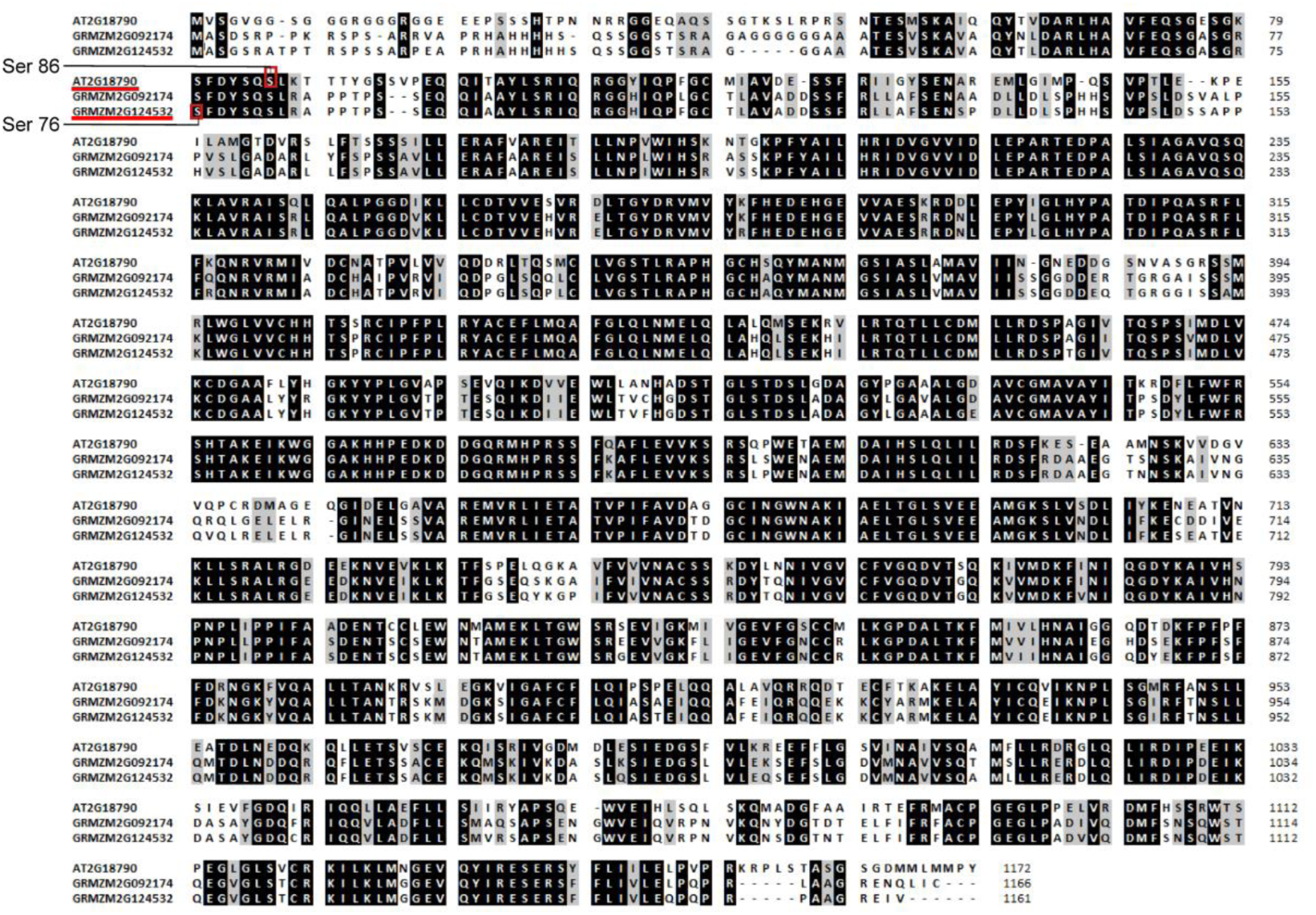
Alignment of the AtPHYB and ZmPHYB proteins The amino acid sequences encoded by AtPHYB (AT2G18790) and two ZmPHYBs (GRMZM2G092174 and GRMZM2G124532) were aligned. The “S” residues outlined in red boxes are the phosphorylation sites Ser86 identified in *Arabidopsis* and Ser76 identified in *Zea mays*.

**Supplementary Figure 6.**
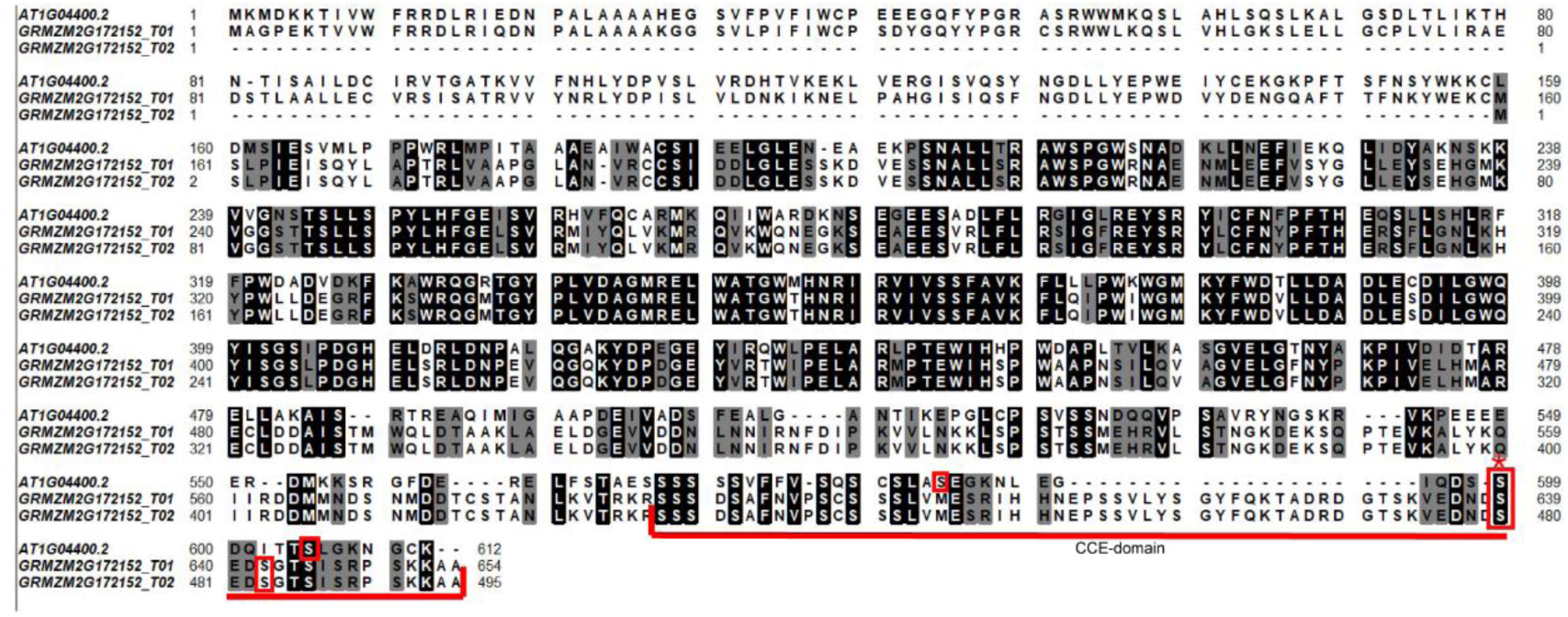
Alignment of the AtCRY2 and ZmCRY2 proteins The amino acid sequences encoded by AtCRY2 (AT1G04400.2) and two ZmCRY2 transcripts (GRMZM2G172152_T01 and GRMZM2G172152_T02) were aligned. The “S” residues outlined in red boxes are the phosphorylation sites identified in *Arabidopsis* and *Zea mays*. The red asterisk indicates the conserved phosphorylation sites.

**Supplementary Figure 7.**
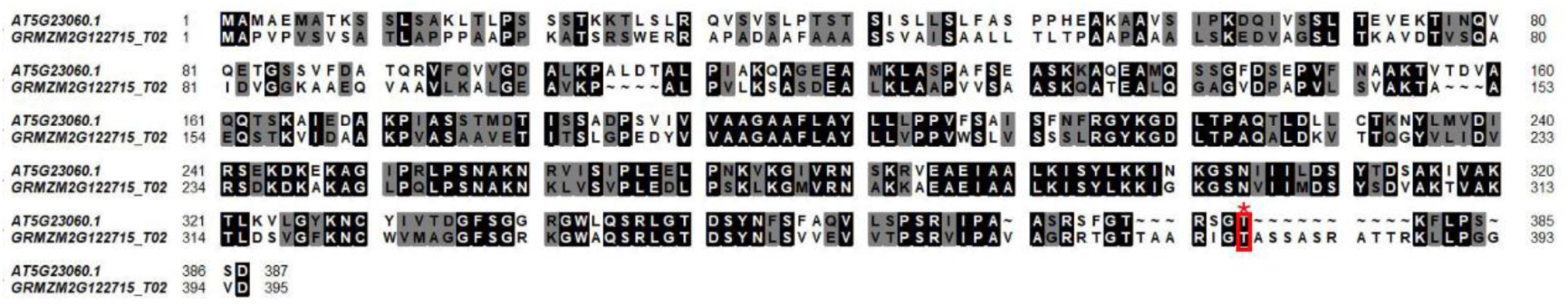
Alignment of the AtCAS and ZmCAS proteins The amino acid sequences of AtCAS (AT5G23060.1) and ZmCAS (GRMZM2G122715_T02) were aligned. The “T” outlined by a red box with asterisk is the phosphorylation site identified in both the *Arabidopsis* and *Zea mays* proteins.

**Supplementary Figure 8.**
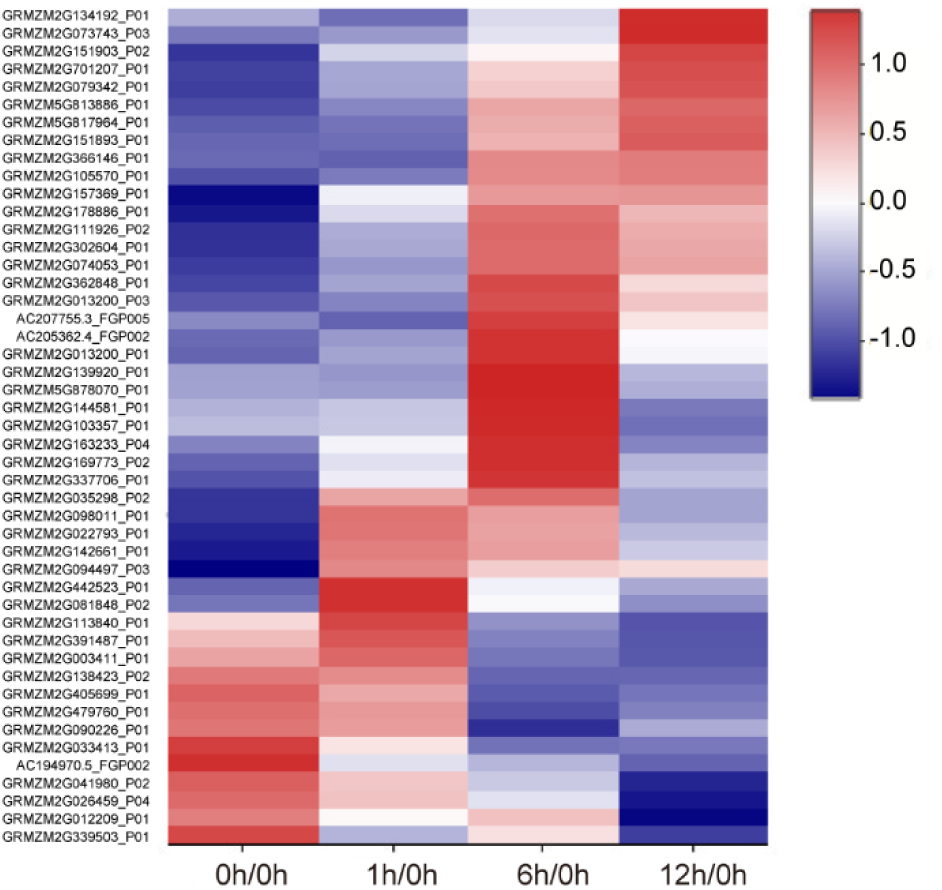
The dynamics of transporters with significant changes in protein abundance during de-etiolation Heat map showing the hierarchical clustering of transporters with significant changes in abundance. The blue color represents low abundance while the red color represents high abundance.

